# ChimeraTE: A pipeline to detect chimeric transcripts derived from genes and transposable elements

**DOI:** 10.1101/2022.09.05.505575

**Authors:** Daniel S. Oliveira, Marie Fablet, Anaïs Larue, Agnès Vallier, Claudia M. A. Carareto, Rita Rebollo, Cristina Vieira

## Abstract

Transposable elements (TEs) produce structural variants and are considered an important source of genetic diversity. Notably, TE-gene fusion transcripts, *i.e.,* chimeric transcripts, have been associated with adaptation in several species. However, the identification of these chimeras remains hindered due to the lack of detection tools at a transcriptome-wide scale, and to the reliance on a reference genome, even though different individuals/cells/strains have different TE insertions. Therefore, we developed ChimeraTE, a pipeline that uses paired-end RNA-seq reads to identify chimeric transcripts through two different modes. Mode 1 is the reference-guided approach that employs canonical genome alignment, and Mode 2 identifies chimeras derived from fixed or insertionally polymorphic TEs without any reference genome. We have validated both modes using RNA-seq data from four *Drosophila melanogaster* wild-type strains. We found ∼1.12% of all genes generating chimeric transcripts, most of them from TE-exonized sequences. Approximately ∼23% of all detected chimeras were absent from the reference genome, indicating that TEs belonging to chimeric transcripts may be recent, polymorphic insertions. ChimeraTE is the first pipeline able to automatically uncover chimeric transcripts without a reference genome, consisting of two running Modes that can be used as a tool to investigate the contribution of TEs to transcriptome plasticity.

## INTRODUCTION

Transposable elements (TEs) are mobile DNA sequences that comprise a large fraction of eukaryotic genomes, from 15% in *Drosophila melanogaster* (1), 45% in humans (2), to 85% in maize (3). Many TE copies have lost their ability to transpose as a result of accumulated mutations and recombination throughout evolution (4). Despite their lack of mobility, such ancient TE insertions may still harbor functional protein domains, alternative splice sites, and *cis*-acting regulatory sequences, as transcription factor binding sites (TFBSs) and polyadenylation (PolyA) sites. Therefore, TEs are a major source of genetic diversity, not only due to their mobilization, but also because they donate protein domains to genes (5–8), and regulatory sequences that modify the expression of nearby genes (9–13). The participation of TE-derived sequences in the host biology is a process called domestication or exaptation (14). The ancestral role of the TE sequence can be domesticated into an essential host function, but it can also be modified, and adapted, into a new exapted function that may also be essential for the host species (14).

Chimeric transcripts are RNAs stemming from two sequences of different origins (15). Hereafter we define chimeric transcripts as mature transcripts that have both gene and TE-derived sequences. These transcripts can be divided into three types: (1) TE-initiated transcripts: chimeric transcripts with a TE transcription start site (TSS) (16, 17); (2) TE-exonized transcripts: TE sequences are incorporated into the transcript either partially or as full-length exons (18–20); and (3) TE-terminated transcripts: chimeric transcripts with a TE transcription termination site (21, 22). TE-initiated and TE-terminated transcripts might modulate gene expression levels either by the presence of TFBSs, PolyA sites, or chromatin changes; while TE-exonized transcripts may alter the protein sequence of coding genes and have a direct effect on the protein function. Regardless of the TE position, such events of TE exaptation and domestication have been associated with many biological roles and are widespread among eukaryotic species (14). In *D. melanogaster*, the *CHKov1* gene produces a chimeric transcript with a truncated mRNA resulting in resistance to insecticide and viral infection (23). In humans, the *SETMAR* gene produces a chimeric transcript containing a *Hsmar1* copy, involved in non-homologous end-joining DNA repair (24). In cancer, TEs become active due to a global hypomethylation state (25) and such activation may generate new chimeric transcripts with detrimental outcomes (26), a process called onco-exaptation (9). For example, in large B-cell lymphoma, the *FABP7* gene has an endogenous retrovirus LTR co-opted as a promoter, generating a novel protein involved in abnormal cell proliferation (27). Recently, such phenomenon has been observed in a larger scale. A pan-cancer study revealed 1,068 tumor-specific TE-initiated transcripts from genes coding antigens, showing the high prevalence of TE exaptation in cancer (28). Therefore, chimeric transcripts have a large impact on host biology, but their study remains hindered by the ubiquitous repetitive nature of TE copies, and lack of methods to identify chimeric transcripts.

Previous studies with Cap Analysis Gene Expression (CAGE) revealed a significant percentage of genes producing TE-initiated transcripts, ranging from 3-14% in humans and mice, depending on the tissue (29). More specifically, in human pluripotent stem cells, chimeric transcripts comprise 26% of coding and 65% of noncoding transcripts (30). In *D. melanogaster*, a study using expressed sequence tags (ESTs) showed that the proportion of genes with chimeric transcripts is only ∼1% (31). Another study used an approach based on cap-enriched mRNA sequencing but with a preliminary transcript elongation step allowing not only to detect TSSs but also to estimate mRNA expression rates (RAMPAGE) (32). This study detected TE-initiated chimeric transcripts in 36 stages of *D. melanogaster* life cycle, and observed that up to 1.6% of all transcripts were chimeras, representing up to ∼1% of all genes (33). More recently, a tissue-specific study has shown that 264 genes produce chimeric transcripts in the midbrain of *D. melanogaster*, corresponding to ∼1.5% of all genes (34).

Several bioinformatic methods have been developed to take advantage of RNA sequencing (RNA-seq) to identify chimeric transcripts, such as CLIFinder (35) and LIONS (36). The former is designed to identify chimeric transcripts derived from long interspersed nuclear elements (LINEs) in the human genome, whereas the latter identifies only TE-initiated transcripts. Both methods need a reference genome and they only detect chimeric transcripts derived from TE insertions present in the reference genome. Therefore, it is impossible to identify chimeric transcripts derived from polymorphic TE insertions that may exist in other populations, strains, or individuals. Finally, the latest addition to chimeric transcript detection, TEchim (34), can detect chimeras with TEs that are polymorphic and absent from the reference genome of *D. melanogaster*, but it is not a pipeline designed to run automatically with any other genome.

In this study, we have developed ChimeraTE, a pipeline that uses paired-end RNA-seq reads to identify chimeric transcripts. The pipeline has two Modes: Mode 1 can predict chimeric transcripts through genome alignment, whereas Mode 2 performs chimeric transcript searches without a reference genome, being able to identify chimeras derived from fixed or polymorphic TE insertions. To benchmark the pipeline, we have used RNA-seq from ovaries of four *D. melanogaster* wild-type strains, for which we have assembled and annotated genomes. We found that ∼1.12% of genes have chimeric transcripts in the ovarian transcriptome, of which ∼88.97% are TE-exonized transcripts. Our results also reveal that the retrotransposon *roo* is the most frequent exonized TE family. In addition, with Mode 2, we found 11 polymorphic chimeric transcripts deriving from TE insertions that are absent from the *D. melanogaster* reference genome. Therefore, this work provides a new strategy to identify chimeric transcripts with or without the reference genome, in a transcriptome-wide manner.

## MATERIAL AND METHODS

### ChimeraTE: the pipeline

ChimeraTE was developed to detect chimeric transcripts with paired-end RNA-seq reads. It is developed in *python3 v.3.6.15* language, and it is able to fully automate the process in only one command line. The pipeline has two detection Modes: (1) genome-guided, the reference genome is provided and chimeric transcripts are detected aligning reads against it; and (2) genome-blind, the reference genome is not provided and chimeric transcripts are predicted for fixed or polymorphic TEs. These Modes have distinct approaches that may be used for different purposes. In Mode 1, chimeric transcripts will be detected considering the genomic location of TE insertions and exons. Chimeras from this Mode can be classified as TE-initiated transcripts (TE located upstream of the gene), TE-exonized transcripts (TE within introns or embedded in gene exons), and TE-terminated transcripts (TE located downstream of the gene). In addition, results from Mode 1 can be visualized in a genome browser, which allows a manual curation of chimeric transcripts in the reference genome. Mode 1 does not detect chimeric transcripts derived from TE insertions absent from the provided reference genome. Mode 2 predicts chimeric transcripts considering the alignment of reads against transcripts and TE insertions, in addition to a transcriptome assembly (user optional). Hence, Mode 2 detects chimeric transcripts from *de novo* TE insertions and an assembled genome is not necessary. In this Mode, two alignments are performed: (1) transcript alignment and (2) TE alignment. Then, based on both alignments, the pipeline identifies chimeric reads that support chimeric transcripts, regardless of the TE genomic location. In Mode 2, since there is no alignment against an annotated genome, it is not possible to classify chimeric transcripts considering the TE position as in Mode 1.

Both ChimeraTE Modes use chimeric reads, which are defined by paired-end reads stemming from TE to exon sequences, as evidence of chimeric transcripts. This method has been widely demonstrated by other authors as a potential source of artifactual reads, mainly due to the occurrence of mixed clusters (*index hopping*) on the Illumina’s flow cell that may be too close to each other, as well as jumping PCR (*in vitro* crossover artifact), generating read pairs that are connecting two cDNA portions that are not joined in the sample (37–40). Indeed, it has been shown that up to 1.56% of all reads produced by Illumina multiplexed approaches might generate chimeric reads (41), including cases that may support chimeric transcripts derived from different genes. These artifactual reads originate more likely from highly expressed genes since there are more molecules on the Illumina’s cell. Conversely, because TE-derived sequences might comprise a low proportion of the transcriptome, artifactual reads from TEs should be produced at a low frequency. Furthermore, it is unlikely that artifactual reads from the same gene and TE family among RNA-seq replicates would be produced. Nonetheless, to avoid including false chimeric reads, both Modes of ChimeraTE only call chimeric transcripts that are detected in at least two RNA-seq replicates (user optional for more).

### ChimeraTE Mode 1: genome-guided approach

In ChimeraTE Mode 1, paired-end RNA-seq reads, genome, and its respective gene/TE annotation are used to predict chimeric transcripts (Figure 1A). We highlight the need for robust gene and TE annotations to submit an analysis to Mode 1, since each TE insertion and exon will be used to identify chimeric reads. Firstly, the genome alignment is performed with *STAR v.2.7.6a* with default parameters (Figure 1B), and transcript expression is assessed with *Cufflinks v.2.2.1* (42) for each RNA-seq replicate. We consider FPKM >= 1 as an expressed gene by default, but it can be changed by the user with the *--fpkm* parameter. Only concordant reads (both reads from the pair are aligned) with unique alignment (*samtools -q 255*) are selected and converted to BED format with *samtools v.1.10* (43) and *bedtools v.2.30.0* (44), implemented in *python3* with *pybedtools v.0.9.0* (45). Split reads are also considered as chimeric reads when one mate of the pair stems from the TE to the exon, or vice-versa (Sup. Figure 1). The alignment position of these reads are considered independently with *bedtools bamtobed-split*. Finally, reads aligned into the forward and reverse strands are separated with *samtools* (43).

**Figure 1:**
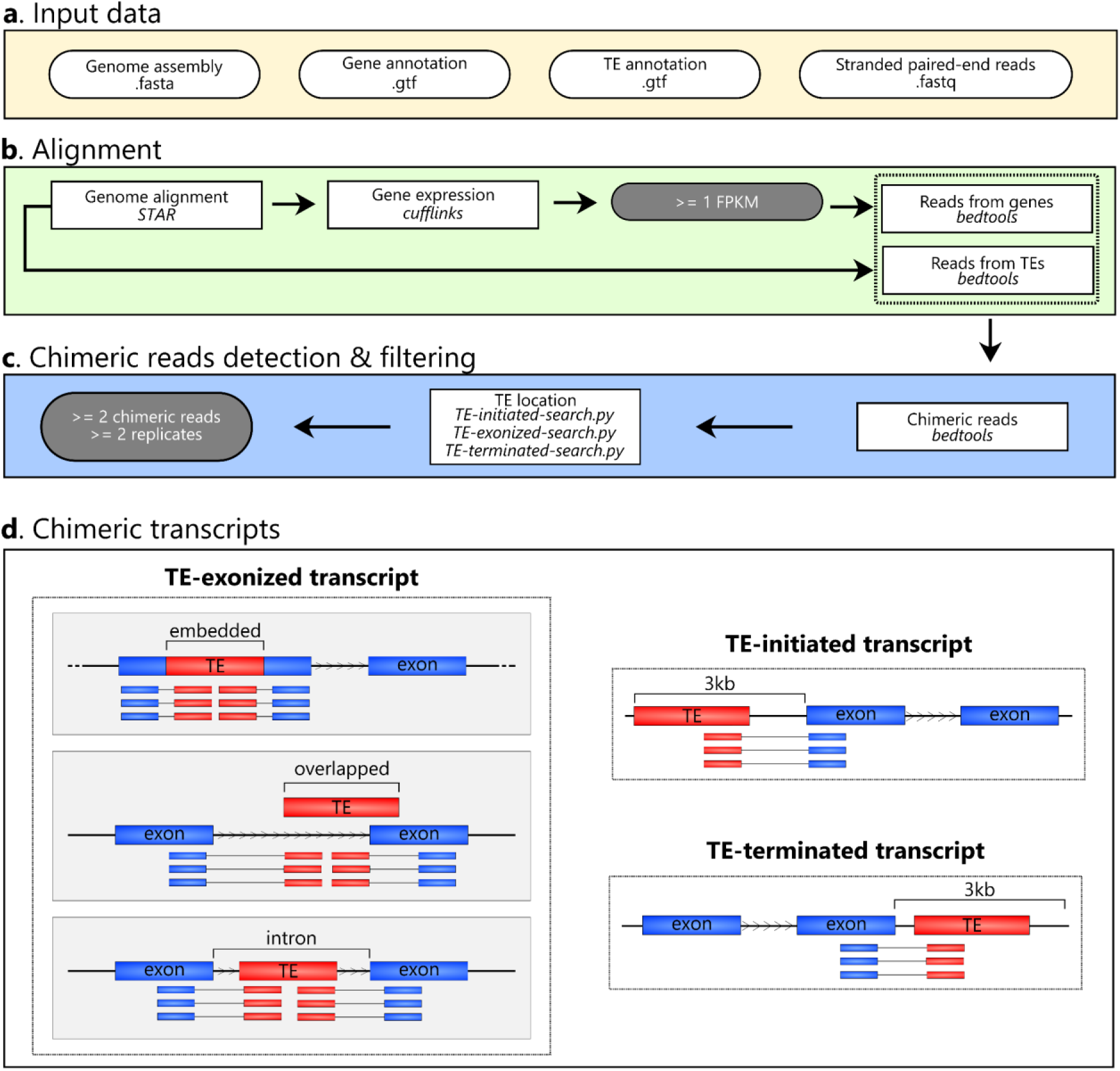
ChimeraTE Mode 1 (genome-guided) workflow. Round white boxes: input data; square boxes: pipeline step; round gray boxes: thresholds that can be modified. **A)** Input data: fasta file with the genome assembly, gtf files with gene and TE annotations, as well as stranded paired-end reads from RNA-seq (fastq). **B)** Alignment: genome alignment is used to calculate gene expression levels. Genes with FPKM =< 1 are removed from downstream analyses. A subsequent list of reads that have aligned against genes or TE insertions is created. **C)** Chimeric read detection & filtering: both read lists are then compared and read pairs that have common reads between the two lists are named chimeric reads, *i.e.*, paired-end reads mapping to a gene and a TE copy. The average of these reads between replicates is used as chimeric read coverage for each putative chimeric transcript. All putative chimeras are then processed with three ChimeraTE scripts to categorize them into TE-initiated, TE-exonized, and TE-terminated transcripts. These steps are performed for all RNA-seq replicates. Finally, all chimeric transcripts present in at least 2 replicates and with at least 2 chimeric reads on average between replicates are maintained. **D)** Chimeric transcripts: Three predictions obtained from Mode 1. Blue boxes: exons; red boxes: TEs; arrowhead in between TE and exon boxes: transcription sense; blue and red boxes linked by a line: chimeric reads. The ChimeraTE mode 1 output is divided into three predictions: (1) TE-initiated transcript: the TE insertion is located upstream of the gene region; (2) TE-exonized transcript: the TE insertion is present within exons (embedded), overlapping exons (overlapped), or introns (intronic); (3) TE-terminated transcript: the TE insertion is located downstream of the gene region.

The IDs from reads that have aligned against exons are identified with *bedtools intersect* (Figure 1B). Then, reads with at least 50% of their length (*--overlap 0.50*) aligned against reference TE copies have their IDs selected (Figure 1B), and TE copies without aligned reads are removed from the downstream analysis. These two lists of IDs are intersected between each other and reads that have aligned to both exons and TEs are identified, generating a raw list of chimeric reads (Figure 1C). TE-initiated transcripts are the first to be searched by ChimeraTE Mode 1. Expressed genes that have TE copies within their 3 kb upstream region (default but adjustable with *--window* parameter) are identified with *bedtools* (44). For each gene, the pipeline will check whether the TE located upstream has chimeric reads shared with the gene exons (Figure 1D). If there are no chimeric reads, both gene and TE are discarded. The same method is applied to identify TE-terminated transcripts, but with genes harbouring downstream TEs up to 3 kb (user adjustable value) (Figure 1D). Finally, TE-exonized transcripts are also identified thanks to chimeric reads aligned to exons and TEs located within genes. Based on the TE position towards exons, they are classified as embedded, intronic and overlapped. Embedded TEs are located entirely within an exon. In these cases, reads aligned only to the TE sequence are not considered chimeric reads (Sup. Figure 1). Indeed, the exon can be artificially divided into two portions, and chimeric reads must have a read (or a split read) aligned to the TE, whereas its mate (or the extension from the split read) aligned to at least one exon portion (Sup. Figure 1). This method avoids autonomous TE expression as evidence of chimeric transcript expression. The same is performed to identify TE-exonized transcripts derived from TEs that are partially overlapping exons but are also located in introns (Sup. Figure 1). Finally, TE-exonized transcripts with TEs located in introns are identified with *bedtools intersect*, and chimeric reads are detected to support chimeric transcripts.

These steps are repeated for all RNA-seq replicates provided in the input. Then, the raw results from replicates are compared, and all chimeric transcripts that were found in >= 2 replicates and have been identified with at least >= 2 chimeric reads on average between replicates are considered true chimeric transcripts (Figure 1C). These thresholds may be changed by the user with *--cutoff* and *--replicate* parameters. Mode 1 output is a table with a list of predicted chimeric transcripts categorized by TE position, with gene ID, TE family, chimeric read coverage, TE location, gene location, and gene expression (Figure 1D).

### ChimeraTE Mode 2: a genome-blind approach to uncover chimeric transcripts

ChimeraTE Mode 2 is the genome-blind approach of the pipeline. The input data are stranded RNA-seq reads, reference TE insertions, and gene transcripts (Figure 2A). The data will be used to perform two alignments with *bowtie2 v.2.4.5* (46), one against all transcripts and another against all TE sequences, both with parameters: -*D 20 -R 3 -N 1 -L 20 -i S,1,0.50* (Figure 2A). To avoid very low-expressed transcripts predicted as chimeras and to ensure reasonable processing time, the SAM alignment is converted to BAM with *samtools v.1.10* (43), and FPKMs are computed for the reference transcripts provided in the input using *eXpress v.1.5.1* (47). Then, all genes with average FPKM < 1 are removed from the downstream analysis (Figure 2B). To identify chimeric reads between TEs and gene transcripts, both alignments are converted to BED with *bedtools v.2.30.0* (44). Among all aligned paired-end reads, the pipeline considers as chimeric transcripts the ones that have at least one read aligned to the TE sequence (singleton mapping) and its mate to the gene transcript, or when both reads have aligned (concordant mapping) to the TE and gene transcript. In order to identify these reads, the TE alignment output is used to create a list with all read 1 IDs that have aligned against TEs, and another list with all read 2 IDs, regardless if their mates have also aligned (concordant mappings or singleton mappings). The same lists are created by using the transcript BED file: (1) read 1 IDs of transcript mapping reads and (2) mate 2 IDs of all mate 2 reads, regardless of mate mapping. All mate 2 IDs that have a TE-aligned read 1 are searched in the list of transcript-aligned mate 2. The same is performed in the opposite direction (TE-aligned read 2, transcript-aligned mate 1). These read pairs will therefore be comprised of two mates from the same pair that were singletons in the alignments, *i.e.*, pairs comprised of one read that has aligned against a TE, and its mate against a gene transcript. The cases in which the chimeric transcript does not have the TE insertion in the reference transcript will be supported only by these singleton chimeric reads (Figure 2D). For cases in which the TE insertion is present inside the reference transcript, chimeric reads supporting it may either be singleton or concordant reads. Therefore, chimeric reads can be concordant reads in both alignments (TEs and genes), or they may be concordant only in the gene transcript alignment and singleton in the TE alignment. Due to the repetitive nature of TEs, short-read alignment methods provide very few unique aligned reads against loci-specific TE copies as most reads align ambiguously between similar TE insertions. Therefore, when a chimeric transcript has been identified involving more than one TE family, the TE family with the highest coverage of chimeric reads is maintained. Subsequently, ChimeraTE uses two chimeric reads as a threshold for calling a chimeric transcript, that can be modified by the user with the *--cutoff* parameter. Such value does not represent transcript expression nor TE expression, but it represents the coverage supporting the junction between a gene transcript (CDSs/UTRs) and a TE sequence. Finally, the output tables show the list of genes and the respective TE families detected as chimeras, reference transcript ID, and the total coverage of chimeric reads supporting it.

**Figure 2:**
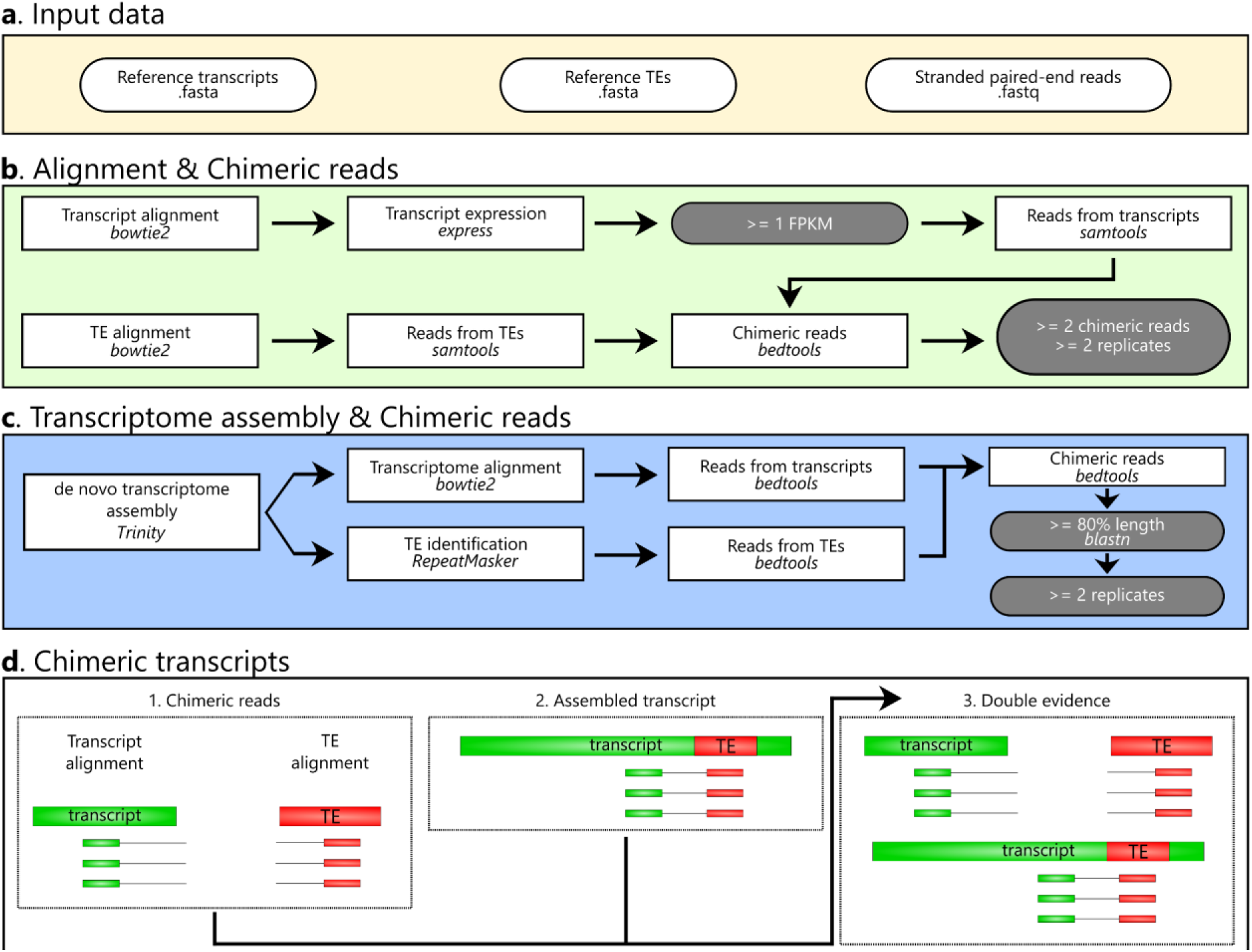
ChimeraTE Mode 2 (genome-blinded) workflow. Round white boxes: input data; square boxes: pipeline step; round gray boxes: thresholds that can be modified**. A**) Input data: two fasta files containing reference transcripts and TE insertions, as well as stranded paired-end reads from RNA-seq (fastq). **B**) Alignment and chimeric reads: The alignment against transcripts is performed and their expression is calculated. Transcripts with FPKM < 1 are removed from the downstream analysis. Next, a list of reads aligned against transcripts is created. Through the alignment of reads against TE insertions, a second list with reads stemming from TEs is also created. Then, mapped paired-end reads and singletons are identified, generating the list of chimeric reads, for all replicates. All chimeric transcripts that have an average of chimeric reads >= 2 and are present in >= 2 replicates are maintained as true chimeras. **C**) Transcriptome assembly and chimeric reads: The *de novo* transcriptome assembly is a non-default option of ChimeraTE Mode 2. It performs a transcriptome assembly and aligns reads against the assembled transcripts. Then, TE insertions in the assembled transcripts are identified with RepeatMasker and the TE reads are recovered. Using the two lists of reads (transcripts and TEs), the chimeric read list is generated and the putative assembled chimeric transcripts are predicted. Next, a blastn is performed between these transcripts and the reference transcripts provided in the input. All transcripts with length >= 80% are selected. The process is repeated for all RNA-seq replicates and chimeric transcripts assembled >= 2 replicates are maintained as true chimeras. **D**) Chimeric transcripts: if the assembly is activated, ChimeraTE mode 2 provides three outputs: (1) Chimeric reads: predicted only based on the method depicted in -B-; (2) Assembled transcripts: predicted only based on the transcriptome assembly method depicted in –C-; and (3) Double evidence: predicted by both methods -B and C-.

The support of chimeric transcripts performed by ChimeraTE Mode 2 is from chimeric reads aligned by an *end-to-end* approach. Such alignment may reduce alignment sensitivity, since exon/TE junctions may be covered by split reads. To mitigate the loss of detection power due to the alignment method with Mode 2, alongside with chimeric read detection using alignments against transcripts and TEs, there is an option to run Mode 2 with a transcriptome assembly approach, which can be activated with *--assembly* parameter (Figure 2C). This approach will use RNA-seq reads to perform a *de novo* transcriptome assembly with *Trinity v2.9.1* (48). In order to identify assembled transcripts that may have TE-derived sequences, genome masking is performed with *RepeatMasker v4.1.2* (49), providing *--ref_TEs*, a custom TE library, or pre-defined TE consensus sequences from *Dfam v3.3* (50), according to the taxonomy level, *i.e*.: flies, mouse, humans. Then, RNA-seq reads are aligned with *bowtie2 v.2.4.5* (46) against the assembled transcripts. Subsequently, the alignment is used to identify whether transcripts containing TE-derived sequences have chimeric reads, including split reads. All assembled transcripts with chimeric transcripts are selected as candidate chimeric transcripts. Next, these candidates are submitted to a homology analysis with reference transcripts using *blastn v2.11.0* (51). Finally, all assembled transcripts with masked TEs that have at least 80% of similarity with reference transcripts across 80% of their length (can be modified with -- *min_length* parameter) are considered chimeric transcripts. All these steps are repeated for all RNA-seq replicates provided in the input. Finally, the list of chimeric transcripts obtained from all replicates with the transcriptome assembly approach is compared, and all chimeras that have been identified with at least >= 2 chimeric reads and were found in >= 2 replicates, are considered as true chimeric transcripts. By activating the *--assembly* option in Mode 2, the output table will provide chimeric transcripts that have been predicted based on different evidences (Figure 2D): (1) only based on chimeric reads; (2) only based on transcript assembly; (3) based on chimeric reads and transcript assembly.

### *D. melanogaster* wild-type strains: genome assemblies and RNA-seq

To assess ChimeraTE’s performance, as well as the efficiency in the identification of chimeric transcripts derived from polymorphic TE insertions, we have mined previously available RNA-seq data from ovaries of four *D. melanogaster* wild-type strains (52), two from France, Gotheron (44_56’0”N 04_53’30”E), named dmgoth101 and dmgoth63; and two from Brazil, São José do Rio Preto (20_41’04.3”S 49_21’26.1”W), named dmsj23 and dmsj7. Sequencing was performed on an Illumina Hiseq (125 bp paire-end reads), with two biological replicates. All RNA-seq libraries were trimmed by quality, and adapters were removed with Trimmomatic (53). Each strain had also its genome previously sequenced by Nanopore long reads and assembled (54). The high-quality assemblies allowed us to manually check whether chimeric transcripts predicted by both ChimeraTE Modes have the predicted TE insertions inside/near genes, as well as manually curate the presence of chimeric reads.

### Running ChimeraTE with *D. melanogaster* data

To run ChimeraTE Mode 1 on the available RNA-seq data, we performed gene annotation in the four *D. melanogaster* genomes with *Liftoff v.1.6.3* (55) using *-s 0.5 -s 0.75 -exclude_partial* parameters and the r6.49 *D. melanogaster* genome (*dm6* strain) available in Flybase as reference. TE annotation was performed with *RepeatMasker v4.1.2* (49), with parameters: *-nolow*; -*norna*; *-s;* and *-lib* with the TE sequence library for *D. melanogaster* provided by Bergman’s lab *v.10.2* (https://github.com/bergmanlab/drosophila-transposons/blob/master/current/D_mel_transposon_sequence_set.fa). Small TE insertions with length < 80 bp were discarded due to the lack of robustness to define them as TEs, following the 80-80-80 rule (56). In addition, due to the presence of simple sequence repeats (SSRs) in many TEs from *D. melanogaster*, such as *roo*, *412*, *tirant* (57), several insertions from *RepeatMasker* might be indistinguishable from SSRs if the repeat region comprises a large proportion of the TE sequence (58, 59). Therefore, to increase the robustness of the TE annotation, we identified the presence of SSRs with *TRF v.4.09* (60) across all insertions annotated on each strain, with parameters *2*, *5*, *6*, *75*, *20*, *50*, *500*, *-m*, *-h,* and removed TEs that presented SSRs over more than 50% of their sequences. We used ChimeraTE Mode 1 with default parameters, setting *--strand rf-stranded*. For ChimeraTE Mode 2, we demonstrate its potential in detecting chimeric transcripts derived from TE insertions that are not present in a reference genome, even though the transcript sequences and TE copies provided as input stemmed from the *dm6* reference genome. Therefore, we have used ChimeraTE mode 2 with RNA-seq from the four wild-type strains listed previously, with reference transcripts from *D. melanogaster* (*dm6* strain) available in Flybase r6.49 (http://ftp.flybase.net/genomes/Drosophila_melanogaster/current/fasta/dmel-all-transcript-r6.49.fasta.gz) (61). The TE annotation for the *dm6* genome was assessed with the same protocol used on the four wild-type strains, with *RepeatMasker* (49). We have used default parameters, except by *--assembly*.

### Additional species data

In order to demonstrate that ChimeraTE can be used with other species than *Drosophila* spp., we applied it to *Arabidopsis thaliana*, *Homo sapiens*, and the fish *Poecilia reticulata* (guppy) RNA-seq datasets. The latter was selected as an example of non-model species. The genome and gene annotation of *A. thaliana* was recovered from NCBI (GCF_000001735.14), version TAIR10.1. The TE annotation was assessed with *RepeatMasker v.4.1.2* (49), with *parameters -species arabidopsis -cutoff 225 -nolow -norna -a -s*. We downloaded two replicates of paired-end RNA-seq from *A. thaliana* leaf, available on NCBI (SRR21230172, SRR21230173). Regarding human data, the genome, gene and TE annotations were retrieved from NCBI (62), version GRCh38.p14. Two paired-end RNA-seq replicates from the K562 cell-line were downloaded from NCBI (SRR521457, SRR521461). Finally, *P. reticulata’*s genome and gene annotation were also retrieved from NCBI (GCF_000633615.1), and TE annotation was assessed with *RepeatMasker v.4.1.2*, parameters*: -species Actinopteri -cutoff 225 -nolow -norna -a -s*. Two RNA-seq replicates from ovary follicular tissue were downloaded from NCBI (SRR17332506, SRR17332508). Both ChimeraTE Mode 1 and Mode 2 were used with default parameters, except by *--threads 32*, and *--ram 64*.

### Benchmarking polymorphic chimeric transcripts with Nanopore genomes

Once chimeric transcripts were identified, we used the high-quality Nanopore assemblies for dmgoth101, dmgoth63, dmsj23, and dmsj7 previously published (54) to confirm whether genes predicted by Mode 1 as chimeric transcripts have indeed the respective TE insertion located near or within them. To do so, we have used an *ad-hoc* bash script to create three bed files from genes: 3 kb upstream; 3 kb downstream, and gene region. Then, we used *bedtools intersect* (44) to identify genes with TEs located in the three regions. For Mode 1 we have randomly sampled 100 chimeric transcripts of each wild-type strain to visualize the alignments performed by Mode 1 on the *IGV* genome browser (63). For Mode 2, all genes not found by *bedtools intersect* with the predicted TE insertion were visualized in *IGV*. In both manual curations, we considered false positives those cases in which we did not find the TE insertion, or we found the TE insertion, but without chimeric reads.

In order to assess the number of chimeric transcripts found by Mode 2 in wild-type strains derived from TE insertions absent in the *dm6* genome, we also used the *ad-hoc* bash script to create the bed files with 3 kb +/- and the gene regions for *dm6*. Then, we used *bedtools intersect* (44) with TE annotation and the gene regions. By using this method, we generated a list of genes with TEs located 3 kb upstream, inside genes (introns/exons), and 3 kb downstream for the *dm6* genome. Then, the polymorphic chimeric transcripts were identified with the comparison of genes with TEs inside/nearby in *dm6* and the list of chimeric transcripts in the wild-type strains. In addition, all chimeric transcripts derived from TEs that were not found in *dm6* were manually curated with the IGV genome browser (63).

### Validation of chimeric transcripts by RT-PCR

*D. melanogaster* strains were kept at 24°C in standard laboratory conditions on cornmeal– sugar–yeast–agar medium. For sampling, 3-7 day old *D. melanogaster* females were immersed in Phosphate-buffered saline solution (PBS) and dissected under a stereomicroscope. Three biological samples were collected in buffer TA (35 mM Tris/HCl, 25 mM KCl, 10 mM MgCl2, 250 mM sucrose, pH 7.5), each composed of 30 pairs of ovaries. All samples were collected on ice in 1.5 mL RNAse free tubes and stored at -80°C until use. Total RNA was extracted using QIAGEN AllPrep DNA/RNA Micro extraction kit (Qiagen 80284) following the manufactor’s guideline. cDNA synthesis was performed on 1 µg of total RNA or water (RT negative control) with BIORAD iScript cDNA systhesis (Biorad 1708891). Primers were designed considering the TE and the exon location with aligned chimeric reads (Sup. Table 1) and PCRs were performed with Taq’Ozyme (Ozya001-1000).

### Sequence and protein analysis of TE-exonized elements

The sequences of TE-exonized elements were extracted from wild-type genomes with *bedtools getfasta* (44), parameter *-s,* using BED files created by ChimeraTE Mode 1. Since the TE reading frame incorporated into the gene transcript is unknown, we recovered all open reading frames (ORFs) with *EMBOSS v.6.6.0 getorf* (64) in the three coding frames, from both strands. Next, the protein domains in these sequences were assessed with *pfam-scan v.1.6* (65), using *Pfam-A.hmm* database from InterPro (66), with default parameters. We applied a filtering step for protein domains with multiple overlapped matches in the same TE insertion, keeping in the downstream analysis only the longest match. We also removed protein domains that had no association with any TE function, following the description from InterPro. Finally, to analyze whether *roo* elements generating TE-exonized embedded transcripts provided a specific motif to the chimeric exons, we extracted their sequences with *bedtools getfasta*, and aligned with *roo* consensus sequence with *MUSCLE v.5.1* (67). Alignment plots were performed with *MIToS v.2.11.1* (68), using the package *Plots*.

### Differential expression analysis

The read count for differential gene expression analysis for the four wild-type strains was obtained from Fablet et al., 2022 (52). The data were normalized with *DEseq2 v.3.10* (69), with a designed model (∼genotype), assuming that the only variable to identify differentially expressed genes between strains must be the genotype. We performed pairwise comparisons with the four wild-type strains, considering differentially expressed genes the ones with adj. p-value < 0.05.

## RESULTS AND DISCUSSION

ChimeraTE predicts chimeric transcripts derived from genes and TEs using two different strategies. Mode 1 is a genome-guided approach that will predict chimeric transcripts from paired-end RNA-seq through chimeric read pair detection. The main advantages of Mode 1 in comparison to Mode 2 are that the first one can detect split reads between exons and TEs, capture chimeric transcripts with low coverage/expression and classify chimeric transcripts according to the TE position: TE-initiated, TE-exonized, TE-terminated transcripts. However, Mode 1 misses chimeric transcripts derived from TE insertions that are absent from a reference genome. Mode 2, which is a genome-blind approach, performs two alignments against reference transcripts and TE copies and then, similar to Mode 1, predicts chimeric reads between these two alignments. In addition, Mode 2 can optionally perform a *de novo* transcriptome assembly able to detect chimeric transcripts and chimeric reads through split read alignment, improving the sensitivity.

### Setting up the datasets for ChimeraTE

Each ChimeraTE Mode requires different input datasets. To run Mode 1 (genome-guided), gene and TE annotations, along with a genome fasta file are necessary. We took advantage of available paired-end RNA-seq datasets from the ovaries of four wild-type strains of *D. melanogaster* (52), for which high-quality genome assemblies were also available (54). We performed gene annotation in the new assemblies using *D. melanogaster*’s genome (*dm6*) from Flybase r6.49 as a reference and obtained ∼17,278 genes per genome (Sup. Table 2). Regarding TE annotation, we found ∼10.48% of TE content in the four wild-type strains, similar to our previous estimations for these strains (54). Filtering out TE insertions smaller than 80 bp, removed ∼5,549 TEs (20.59%) across the four wild-type strains. In addition, to remove potential mapping artifacts, we discarded ∼752 TE copies (2.79%) per strain that harbored SSRs longer than half the TE length. On average across strains, 85% of the TEs filtered out due to the presence of SSRs were *roo* elements (Sup. Figure 2). These *roo* copies were around 112 bp (Sup. Figure 3), which is the same length as the known SSR present in the roo consensus sequence (57). In total, 123 TE families in the four genomes were uncovered, comprising a mean of ∼21,402 TE insertions (*standard deviation =* 883). In all genomes the TE content in bp is higher for LTR, then LINE elements, followed by DNA and Helitron families (Sup. Table 2) (70). In order to run Mode 2 (genome-blind), we used reference transcripts from Flybase r6.49 and performed TE annotation on the reference *dm6* genome. We have obtained 29,086 TE insertions, representing 16.37% of the genome content, following similar TE family proportions as seen for the four wild-type genomes (Sup. Table 2). Nevertheless, the *dm6* genome TE annotation shows an extra ∼6% of TE content compared to the wild-type genomes, mainly due to LTRs and LINEs (Sup. Table 2). The *D. melanogaster’s* heterochromatin corresponds to ∼32.5% of its genome (71), and includes repeat sequences, as TE copies (72). The wild-type assemblies have a high coverage for euchromatin regions, since 96% of gene content was recovered, but a poor heterochromatin coverage, corresponding to the lack of TEs compared to the *dm6* genome. However, the search for chimeric transcripts with ChimeraTE Mode 1 is restricted to the gene region and its 3 Kb boundaries, hence, the lack of heterochromatic TE insertions does not impact the chimeric transcript search.

### ChimeraTE Mode 1 reveals that 1.12% of genes produce chimeric transcripts in *D. melanogaster* wild-type strains

ChimeraTE mode 1 was run on the four wild-type strain genomes and their respective ovarian RNA-seq data (52). Across all strains, we found 327 genes producing 408 chimeric transcripts, accounting for 1.83% of the total genes in *D. melanogaster*, 4.04% of the expressed genes (FPKM >1), and representing 1.12% of all genes when normalized by the four strains. Previous studies have found similar proportion of genes generating chimeras in *D. melanogaster.* Among them, Coronado-Zamora et al. is the only study that performed chimeric transcripts search in multiple strains and different tissues, including ovaries, for which the authors found 1.41% of genes with chimeric transcripts when normalized by all strains (73). To verify whether ChimeraTE identified the correct TE family for each chimera, we have compared the genomic coordinates of TEs and genes (3 kb upstream/downstream and inside genes) with *bedtools intersect* (44) and predicted genes with TE copies in these regions. All chimeric transcripts in the four wild-type strains had at least one TE insertion in the expected position from the predicted TE family (Sup. Table 3-6). In addition, we randomly selected 100 chimeric transcripts in each wild-type strain to visualize in IGV (63) and confirm the presence of chimeric reads as expected (Figure 3A). All the 400 manually inspected chimeras were correctly found in the genome browser. Among all genes generating chimeric transcripts, around 89% are TE-exonized (61.84% correspond to TE-exonized embedded, 17.18% to TE-exonized intronic, 9.95% to TE-exonized overlapped), 8.64% TE-terminated and 2.36% TE-initiated transcripts (Figure 3B). It is important to note that ChimeraTE does not classify an internal TE copy as either a TE-initiated or a TE-terminated transcript, even if the transcript does indeed start, or end, at the TE sequence. TE-initiated and TE-terminated repeats are necessarily outside of gene regions as *per* ChimeraTE categories (see Methods). Therefore, one should assume that the high prevalence of TE-exonized transcripts (∼89 %) might be associated with potential cases of mis annotated TE-initiated and TE-terminated transcripts. Indeed, a previous study in *D. melanogaster* has shown that ∼2.8% of genes have TE-derived promoters in embryos, pupae, and larvae tissues (33).

**Figure 3:**
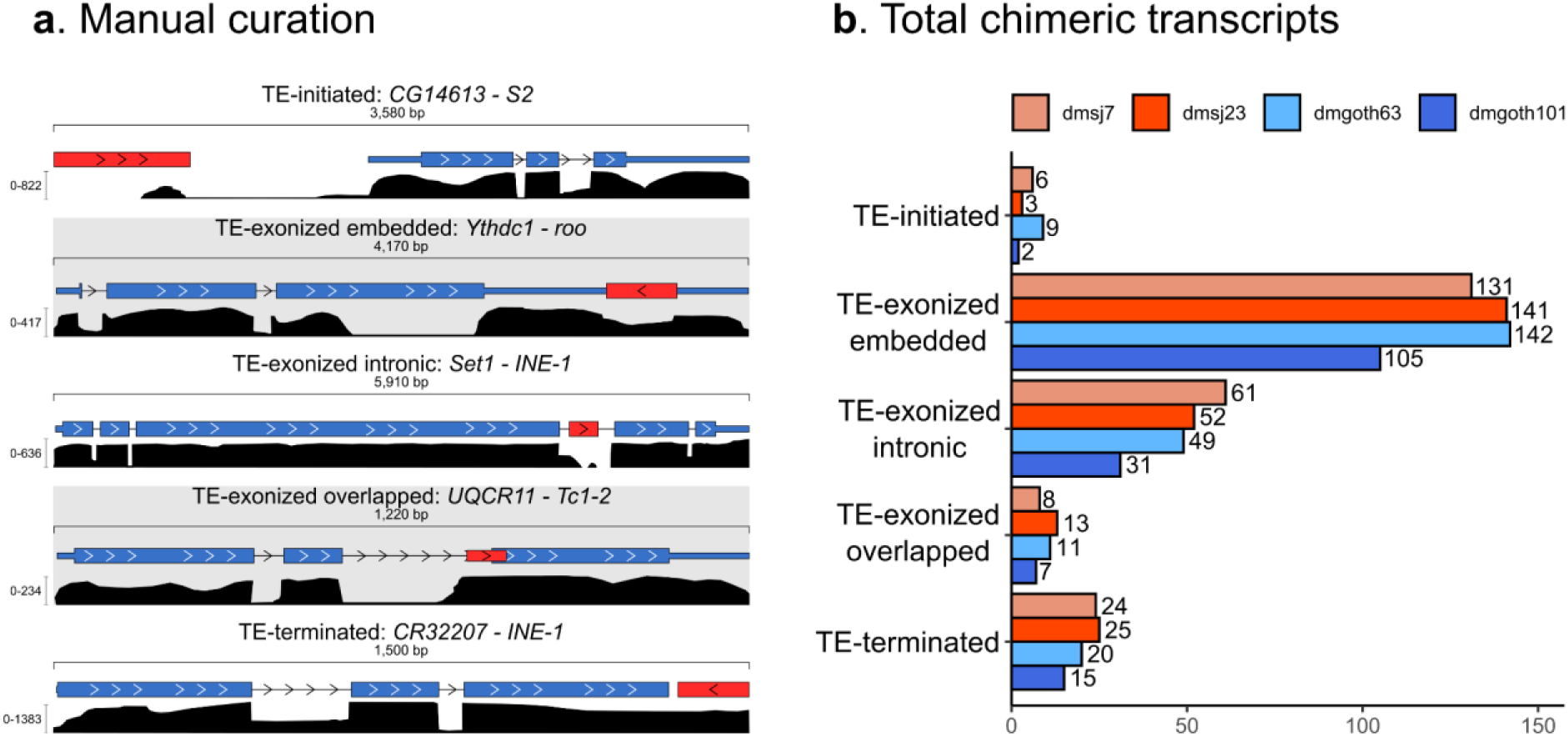
General results from Chimera Mode 1. **A**) Five examples of chimeric transcripts manually curated with the IGV genome browser. Red boxes: TE insertion; blue boxes: exons and UTRs; black density graphs: coverage of RNA-seq reads; head arrows: transcription sense, blue and red boxes linked by a line: chimeric reads. **B**) Total genes generating chimeric transcripts, following the TE position classification in the four wild-type strains.

A recent analysis in *D. melanogaster* ovaries has found 884 chimeric transcripts derived from 549 genes, among five wild-type strains (73), indicating that a gene is capable of producing multiple chimeric transcripts involving different TE copies. In agreement, TE-exonized chimeric genes detected by ChimeraTE, show evidence of multiple TE sequences acting as chimeric transcripts (22.6% of TE-embedded genes, 30.95% for overlapped, and 30.34% for intronic). Nevertheless, TE-initiated transcripts do not involve more than one TE *per* gene. Among 46 TE-terminated transcripts found in all strains, only five genes were reported with more than one TE copy: (1): *NADH dehydrogenase* (*CG40002*) in dmgoth63; (2–3) *Max* and *rudimentary-like* in dmsj23; and (4–5) *six-banded*, *CG14464* in dmsj23. All of them, except *CG14464*, were associated with two TE insertions from the same TE family, located close to each other (Sup. Figure 4), suggesting they might be multiple hits from *RepeatMasker* corresponding to only one TE copy (74). Thus, multiple TE copies producing chimeric transcripts exist only when TE copies are within genes. We then tested whether the number of TEs within genes has a correlation with the presence of multiple TE-exonized chimeras. By using all the genes expressed in ovaries (FPKM > 1) with at least one TE insertion within, we found weak positive correlation between them (Sup. Figure 5), among which the highest was found in dmsj23 (*Pearson; r = 0.22; p = 2.6E-08*), and non-significant for dmgoth101 (*Pearson; r = 0.06; p = 0.08*). However, when we take into account the number of TE copies per TE family present within genes, and the respective number of chimeras generated by TE family, we observed a positive correlation (*Pearson*; *r*=*0.87*; *p* < *2.2E-16*) (Sup. Figure 6). Therefore, TE-rich genes are not the most frequent among the ones generating chimeras, but TE families with high copy number within genes are the ones likely to produce TE-exonized transcripts.

Among all chimeric transcripts predicted by ChimeraTE Mode 1, 11 chimeras have been previously described in *D. melanogaster* using EST data (31). It is important to note that the authors did not consider heterochromatic regions, as well as chimeric transcripts derived from INE-1 elements. From these 11 chimeras, five were found in all strains: *Ssdp* (gene) - *HMS-Beagle* (TE family); *Agpat1*-*1360*; *anne-1360*; *Atf6-Xelement*; and *ctp-HMS-Beagle*. The other six chimeric transcripts are absent from some strains: *CG6191-jockey*; *CG15347-HB*; *CG3164-McClintock; Svil-roo; CHKov1-Doc*; and *Kmn1-pogo* (Sup. Table 7). We manually checked these chimeric transcripts in the IGV genome browser and we confirmed the presence of the TE families predicted by ChimeraTE (Sup. Table 7). In addition, we also found three chimeras reported previously with RAMPAGE: *CG31999*-*Tc1-2*; *CkIIα*-*1366*; *Svil*-*roo* (33). Finally, in another study on *D. melanogaster* midbrain, 264 genes were shown to produce chimeric transcripts using single-cell RNA-seq data (34). The authors demonstrate that retrotransposons have splice donor and acceptor sites that generate new chimeric isoforms through alternative splicing (34). Despite the differences between tissues and methods, we have found 22 chimeric transcripts identified by this study (Sup. Table 7), of which two were previously found in ESTs (31). Taken together, the genome-wide analysis performed by ChimeraTE Mode 1 has uncovered 296 genes with chimeric transcripts in the *D. melanogaster* ovarian tissue for the first time.

### ChimeraTE Mode 2: a method to uncover chimeric transcripts without genome alignment

*D. melanogaster* has a high rate of TE insertion polymorphism across worldwide populations (75–78). To test the ability of ChimeraTE Mode 2 in detecting chimeric transcripts derived from TE insertions absent from a reference genome, we used RNA-seq from the four wild-type strains with transcript and TE annotations from the *D. melanogaster* reference genome (*dm6*). ChimeraTE Mode 2 may predict chimeric transcripts based on two types of evidence: either using chimeric reads only or by taking advantage of *de novo* transcriptome assembly. Chimeric transcripts with both evidences are named “double-evidence” (chimeric reads and transcript assembly - Figure 2B and C). Among the four wild-type strains, ChimeraTE Mode 2 has identified 324 genes (Sup. Table 8-11) producing 339 chimeric transcripts (Figure 4A), representing 1.81% of the total *D. melanogaster* genes, and 1.07% when normalized by the four wild-type strains. Comparing the “chimeric read” approach with the “transcript assembly” one, the former method found ∼51% as many chimeric transcripts than the detection by transcriptome assembly (Figure 4A). This is probably due to the technical limits of *de novo* transcriptome assembly of repetitive-rich transcripts (79), which may lead to the lack of chimeric transcript isoforms. Between both methods, we did not find any correlation with chimeric read coverage and gene expression (FPKM), indicating that even low expressed genes (FPKM < 10) can be detected by both methods independently (Figure 4B). Double evidence chimeras represents ∼14% of all chimeras detected by Mode 2, and they have been found only with high chimeric read coverage (Figure 4B).

**Figure 4:**
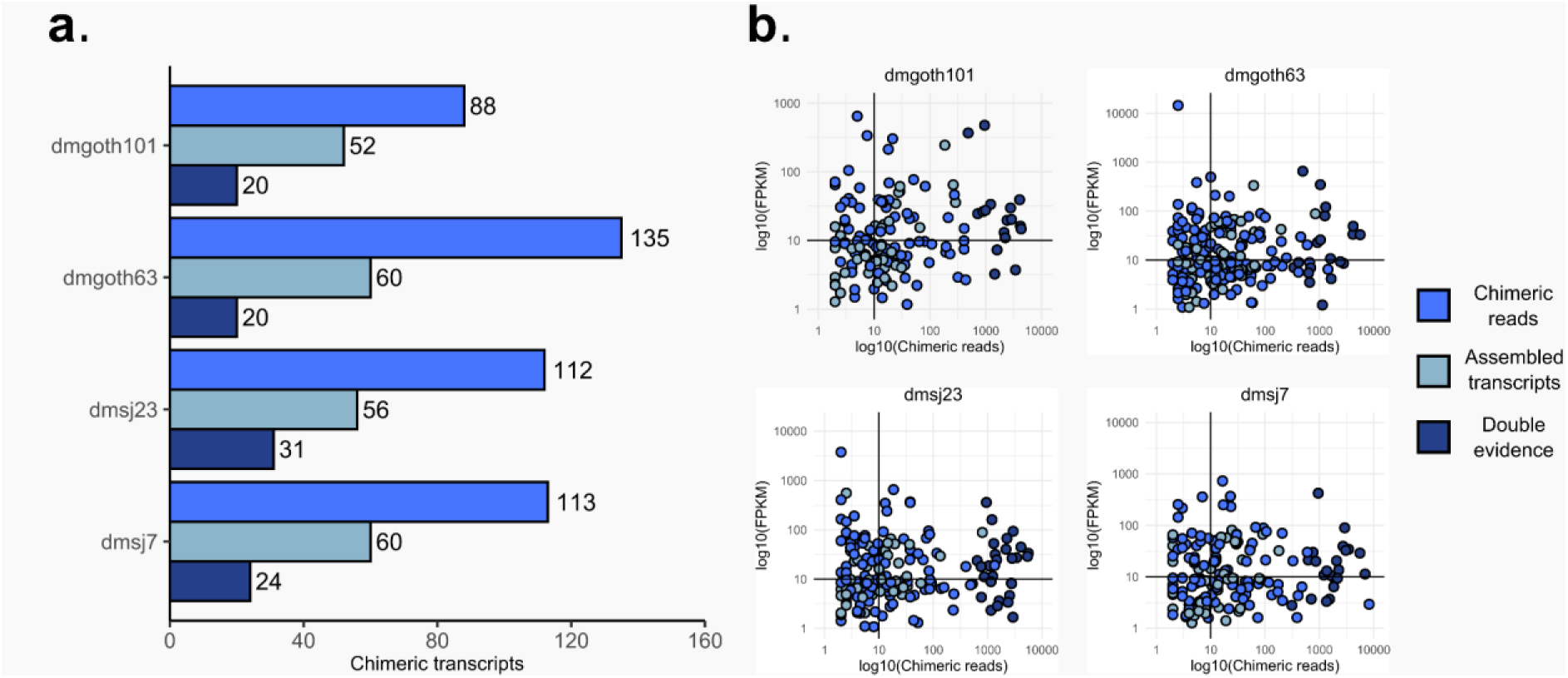
**A**) Total number of chimeric transcripts found by ChimeraTE mode 2. “Assembled transcripts”: chimeric transcripts detected only by the method of transcriptome assembly (Figure 2C). “Chimeric reads”: chimeric transcripts detected only by the method of chimeric reads (Figure 2B). “Double evidence”: chimeric transcripts detected by both methods. **B**) Correlation between chimeric read coverage and gene expression of chimeric transcripts found by Mode 2. In all strains, we did not observe correlation between gene expression (FPKM) and the coverage of chimeric reads between the three categories of evidence. All double evidence chimeras were found with high coverage of chimeric reads, suggesting high reliability of chimeras when both methods (chimeric reads and transcriptome) are considered together.

As each chimeric transcript was detected based on reference transcripts and TE insertions, we used the Nanopore assemblies to manually inspect the presence of the predicted TE family inside and near (+/- 3 kb) genes, with the intersection of genomic coordinates from genes and TEs with *bedtools intersect* (44). We considered true chimeric transcripts the cases in which we found in the assembly the presence of the predicted TE insertion inside/near the gene. The alignment of RNA-seq libraries against the genome sequence was also used to confirm the presence of the TE insertion, as well as the presence of chimeric reads, with the IGV genome browser (63). The manual curation was performed with the three groups of results from Mode 2 (“chimeric reads”, “transcriptome assembly” and “double-evidence”). We found that 96.30% of all chimeric transcripts predicted by double evidence were true, whereas we observed 90.59% from transcriptome assembly and 73.13% from chimeric reads. Therefore, taken together, ChimeraTE Mode 2 has provided a reliable inference with 86.68% sensitivity, based on genomic manual curation.

The main difference of ChimeraTE Mode 2 from the previous pipelines is its ability to detect chimeric transcripts derived from TEs that are absent in a reference genome, using RNA-seq from non-reference individuals/cells/strains. We first identified in the *dm6* reference genome the genes with TEs located upstream, inside, and downstream genes. We found that 2,239 genes in the *dm6* genome have TEs located 3 kb upstream, 1,863 inside (introns and exons), and 2,320 downstream. These genes were selected as candidates to produce chimeras in the *dm6* genome and then compared with the list of chimeric transcripts generated by ChimeraTE Mode 2 in the four wild-type strains. In addition to the comparison with *dm6* genes harboring TEs inside/near, we have manually curated the TE insertions and the presence of chimeric transcripts with the IGV genome browser (63). Such manual curation was performed with the wild-type genomes, by checking the presence of the TE and chimeric read alignments. For instance, the *Mps1-FB* chimera is an interesting case, since it is the only chimeric transcript specific to French populations, as we found it in both dmgoth101 and dmgoth63. In the *dm6* reference genome, *Mps1* has an overlap between its 3’ UTR and the 3’ UTR of the *alt* gene, located on the other strand (Figure 5A). The same distribution of both genes is found in the Brazilian strains, dmsj23, and dmsj7 (Figure 5B). Conversely, in dmgoth101 and dmgoth63, there is a gap of ∼9,500 bp between *Mps1* and *alt*, with four small *FB* copies of ∼412 bp in length (Figure 5C), indicating that it may be an old insertion. Furthermore, one of the *FB* copies is located downstream of the *alt* gene, which has also been identified as TE-terminated downstream in dmgoth63 and dmgoth101. This result shows that ChimeraTE Mode 2 can detect chimeric transcripts derived from TEs that are not present in the reference genome.

**Figure 5:**
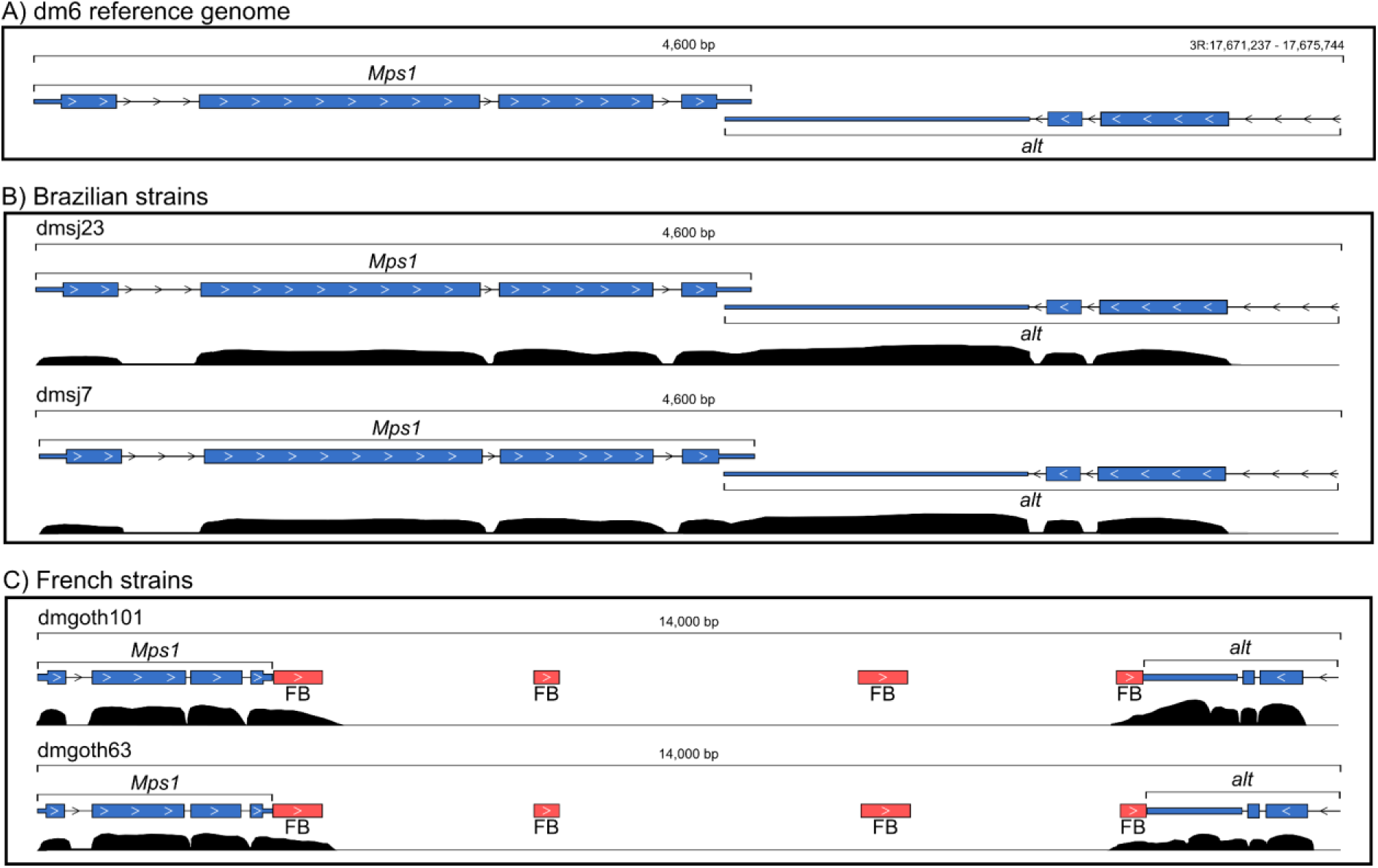
*Mps1* gene and its downstream region in the genomes of the four wild-type strains. **A**) *Mps1* in the *dm6* reference genome and the *alt* gene located downstream to it, on opposite strands and with overlapped 3’ UTRs. **B)** In the dmsj23 and dmsj7, *Mps1* and *alt* are distributed as found in the reference genome. **C**) In dmgoth101 and dmgoth63, there is a *FB* insertion located downstream to *Mps1*, which has chimeric reads supporting a TE-terminated downstream in both strains.

Taking into account all chimeric transcripts detected by Mode 2, we found 11 genes (3.40%) generating chimeric transcripts derived from TEs that are absent from the *dm6* genome (Table 1). There are specific chimeras from French strains: *Mps1-FB*, *CG1358-S,* and *CG46280-POGON1* genes, but only *Mps1* had chimeric transcripts in both French dmgoth63 and dmgoth101 strains, whereas *CG1358* was found as chimera only in the dmgoth101, and *CG46280* only in dmgoth63. The *Ythdc1-roo* chimera was observed with a strain-specific polymorphic *roo* element inside an exon in the dmgoth63 strain. Regarding the Brazilian strains, we found two TE insertions present in both dmsj23 and dmsj7, involving the genes *cic* and *TrpRS-m*, but they were found as chimeras only in the dmsj23 strain. We also found *rb*, *r-1*, and *ArfGAP1* with dmsj23-specific TE insertions, whereas in the dmsj7 we found *caps* and *Ubi-p5E* with specific TE insertions giving rise to chimeric transcripts.

**Table 1:** 11 polymorphic chimeric transcripts from TE insertions that are not present in the *dm6* reference genome. The red triangle in a line represents the presence of a chimeric transcript; the white triangle in a line represents the presence of the TE insertion but without a chimeric transcript; the line without triangles represents the absence of the TE insertion.

| Gene | TE | TE location | Strains |  |  |  |
| --- | --- | --- | --- | --- | --- | --- |
|  |  |  | dmgoth101 | dmgoth63 | dmsj23 | dmsj7 |
| FBgn0000063 - Mps1 | FB4 | TE-terminated | —▲ | —▲ | — | — |
| FBgn0003210 - rb | MDG3 | TE-terminated | — | — | —▲ | — |
| FBgn0003257 - r-l | Stalker2 | TE-terminated | — | — | —▲ | — |
| FBgn0020655 - ArfGAP1 | POGO | TE-terminated | — | — | —▲ | — |
| FBgn0023095 - caps | POGON1 | TE-exonized | — | — | — | —▲ |
| FBgn0027616 - Ythdc1 | ROO | TE-exonized | — | —▲ | — | — |
| FBgn0033196 - CG1358 | S | TE-terminated | —▲ | —△ | — | — |
| FBgn0036763 - TrpRS-m | PROTOP_A | TE-exonized | — | — | —▲ | —△ |
| FBgn0086558 - Ubi-p5E | Gypsy | TE-initiated | — | — | — | —▲ |
| FBgn0262582 - cic | POGON1 | TE-exonized | — | — | —▲ | —△ |
| FBgn0283438 - CG46280 | POGON1 | TE-exonized | —△ | —▲ | — | — |

To evaluate Mode 2 accuracy in detecting chimeric transcripts derived from TEs absent in the reference genome, we have performed RT-PCRs for 10 out of 11 cases predicted (it was not possible to design specific primers for the *TrpRSm-protop_a* chimera). The primers were designed considering the TE and the exon location with aligned chimeric reads (Sup. table 1). For *Mps1*-*FB*, *caps-pogon1*, *Ythdc1*-*roo*, *Ubip5E-gypsy, CG46280*-*pogon1*, the amplified fragment sizes were in agreement with ChimeraTE Mode 2 predictions (Sup. Figure 7A-E). The *r1-stalker2* chimera was amplified in both Brazilian strains, even though the TE insertion generating this chimeric transcript was absent in dmsj7 genome assembly (Sup. Figure 8A). We hypothesized that this might be due to a low-frequency TE in the dmsj7, absent from the assembly and without enough read coverage in the RNA-seq to predict it. Indeed, we confirmed the presence of this TE insertion by PCR in both Brazilian strains (Sup. Figure 8B). The chimeric transcript from *rb-mdg3* was detected only in dmsj23 by ChimeraTE Mode 2, but the RT-PCR assay was successful only in the dmsj7 strain (Sup. Figure 9). The chimera derived from *CG1358-S* was predicted only in dmgoth101 (Sup. Figure 10), despite the TE insertion being also present in dmgoth63 genome. By RT-PCR, we detected this chimeric transcript in both strains, suggesting that it was not detected in dmgoth63 by ChimeraTE due to Illumina coverage issues. A similar result was found for the *cic-pogon1* chimera, for which we amplified the chimeric transcript in both Brazilian strains, but it was detected only in dmsj23 by ChimeraTE (Sup. Figure 11). Altogether, these results indicate that Mode 2 can predict chimeric transcripts from TEs absent from the reference genome, but this prediction depends upon the coverage of chimeric reads in the RNA-seq data.

### Differences between Mode 1 and Mode 2

ChimeraTE Mode 1 and Mode 2 use different alignment strategies and downstream approaches to detect chimeric transcripts (see Methods). In order to test whether these differences can lead to distinct outputs, we compared chimeras from Mode 1 and Mode 2 by using the same RNA-seq libraries from four *D. melanogaster* wild-type strains. Taking all strains together, Mode 2 uncovered 171 chimeric transcripts generated by 165 out of the 327 genes (50.46%) identified by Mode 1, representing ∼25% of all TE-initiated transcripts detected by Mode 1; ∼42% of TE-exonized; ∼29% of TE-terminated (Figure 6A). In addition, we observed 15 genes generating chimeric transcripts in both Modes, but from different TE families. These results indicate that ChimeraTE Mode 2 had low sensitivity (27.39%) to detect chimeric transcripts from TE insertions near genes. In addition, for chimeras derived from TE inside genes, 37.21% of them were detected by the transcriptome assembly approach in Mode 2, showing the relevance of performing this optional analysis. However, it must be considered that *--assembly* performed by Mode 2 is time-consuming, as well as hardware-consuming (Sup. Table 12).

**Figure 6:**
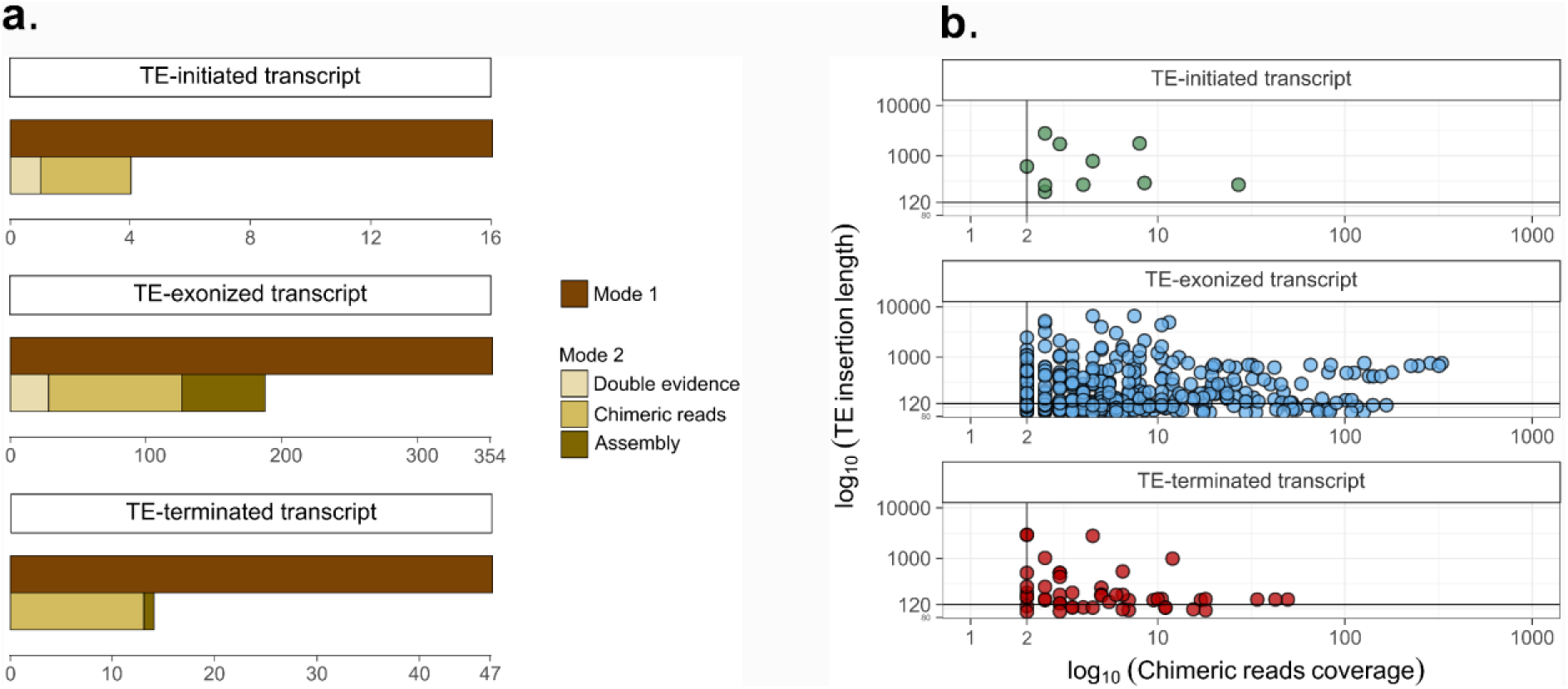
**A**) Total chimeric transcripts detected by ChimeraTE mode 1 and ChimeraTE Mode 2. The three brown boxes depict the type of evidence used by Mode 2 to support the chimeric transcripts (see Methods). Mode 2 had more efficiency to detect chimeric transcripts derived from TEs inside exons than near genes. **B**) Chimeric transcripts found by Mode 1, but not by Mode 2. 47.55% of all chimeric transcripts detected only by Mode 1 have TEs shorter than the read length (120 nt), and 32.51% of chimeras with TEs longer than reads have low chimeric reads coverage (10 chimeric reads). These factors explain the differences between results found by both Modes.

In both ChimeraTE Modes, the main evidence used to detect chimeric transcripts is the presence of chimeric reads, which are paired-end reads spanning between TE and gene sequences. In Mode 1, at least 50% of one read (default parameter) from the read pair must align against the TE insertion, whereas in Mode 2 the whole read must align against a TE copy. Therefore, the alignment method performed by Mode 2 does not allow the detection of chimeric transcripts derived from TE copies shorter than the read size. Hence, we investigated whether chimeric transcripts found with Mode 1, but not with Mode 2, are generated by TE insertions shorter than the sequenced reads. From 237 chimeric transcripts predicted only by Mode 1, we found that 77 (∼32%) have TEs shorter than the read size (120 nt), making it impossible to detect them with Mode 2 (Figure 6B). It is important to highlight that TEs longer than reads may have splice sites, generating chimeric transcripts with a small TE-derived fragment, not being detected by Mode 2 as well. Furthermore, we observed from Mode 2 results that 121 (∼57%) chimeric transcripts from TEs longer than the reads have coverage lower than 10 chimeric reads. We hypothesized that Mode 1 may have substantially more TE-aligned reads than Mode 2, due to the use of strain-specific TE insertions and by counting split read alignment.

ChimeraTE Mode 1 is dependent on a reliable genome annotation for both genes and TEs, contrary to Mode 2. Indeed, we found in total, seven chimeric transcripts detected only with Mode 2 (Sup. Table 13), which are derived from genes that were not annotated in the wild-type genomes. We compared them with the *dm6* genome and we found all seven have the predicted TE family inserted near/inside the gene, reinforcing these annotations (Sup. Figure 12). For instance, the chimeric transcript *CG3164-McClintock* was detected by Mode 2 with double evidence in the four wild-type strains, and it is not annotated in any of the four genomes, perhaps due to its location in the telomeric region of the 2L chromosome (Sup. Figure 12G).

ChimeraTE Mode 1 and Mode 2 detected 327 and 324 genes producing chimeric transcripts respectively, of which 180 were found by both methods. In Mode 2, chimeras with TE-derived sequences smaller than the read length are not detected. However, Mode 2 can detect TEs that are absent from the reference genome, along with low-frequency TE insertions of the wild-type strains at the population level, which are not present in the assembled Nanopore genome. Since low-frequency Nanopore reads are discarded during the genome assembly (54), Mode 1 is not able to detect them, whereas Mode 2 can. Furthermore, Mode 1 detects chimeric transcripts derived from TEs inside or +/- 3 kb from genes. TEs located farther away from genes and involved in chimeric transcripts are not detected. Such chimeras have been reported in *D. melanogaster* (31), as well as TEs acting as distal *cis*-regulatory elements (80, 81). Therefore, although we could consider them as potential false positives all cases in which the TE was not found inside/near a gene in the manual curation, we speculate that part of these cases found only by Mode 2 might be either from low-frequency TEs in the pool of individuals sequenced, or from TEs located far from genes.

### Evaluation of chimeric transcripts in dmgoth101 through Nanopore RNA-seq

Short-read RNA-seq may introduce biases during library preparation (e.g., reverse transcription, second-strand synthesis, PCR amplification) (82), or even during transcriptome assembly, with loss of information regarding isoform diversity, especially for those with low-expression and repetitive regions (83), which might be the case of many chimeric transcripts. In the past years, the third-generation of long-read sequencing has been used to mitigate such biases, because there is no need to perform transcriptome assembly. Thus, to evaluate the chimeric transcripts predicted by ChimeraTE Mode 1, we used long reads produced with Oxford Nanopore Technology (ONT) from dmgoth101 ovaries (84). We did not investigate chimeras from Mode 2 with long reads, due to the lack of information regarding the genomic position of TEs generating them. The long reads were aligned to the dmgoth101 genome and then checked for the presence of all chimeric transcripts detected by ChimeraTE Mode 1 in dmgoth101. The presence of the chimeric transcripts was confirmed when the long read was aligned into the gene exons, and it had the TE-derived sequence as part of the reads. From two TE-initiated transcripts, we found only *CG14613-S2* chimera by ONT reads. In TE-terminated transcripts, six (40%) out of 16 were also observed by ONT data. Finally, in TE-exonized transcripts, we identified 89 (62.24%) out of 143 embedded TEs, 12 (52.17%) out of 23 overlapped TEs, and 13 (46.43%) out of 28 intronic. The undetected chimeras through ONT reads might be associated with the long-read library construction protocol, TeloPrime, which was used to generate full-length cDNA libraries. Such a protocol might miss low expressed isoforms, as well as truncated 3’ ends due to RNA degradation *in vivo* or during RNA manipulation (85). We investigated whether short-read coverage could explain the lack of detection of certain chimeric transcripts with ONT data. Taking all dmgoth101 chimeras from Mode 1, chimeric transcripts found by both ChimeraTE and ONT data had an average chimeric read coverage of 69.44, whereas the undetected ones had 10.83. Therefore, these results indicate that chimeras’ detection with ONT reads is likely to be more efficient with highly expressed chimeric transcripts. Overall, our results demonstrate that 50.16% of all chimeric transcripts detected in the ovary transcriptome of *D. melanogaster* with ChimeraTE Mode 1 can also be found by ONT RNA-seq.

### Replicability and coverage of chimeric transcripts in both ChimeraTE modes

The ability to detect TE expression is an important factor to identify chimeric reads. The different strategies of alignment to identify chimeric reads in Mode 1 and Mode 2 cause differences in the sensitivity of chimeric transcript detection (Figure 6A). We investigated whether such differences might be associated with the detection of TE-derived reads. We found that Mode 1 is more efficient than Mode 2 in detecting reads aligned against TE insertions (Sup. Figure 13), as well as for chimeric read detection (Figure 7A). Indeed, the proportion of chimeric reads in Mode 1 and Mode 2 is 0.34% and 0.04% of the total library sizes, respectively. The power of chimeric read detection from the two Modes is different because of the alignment strategies used. Despite the differences, both modes show significant positive correlations between the library size and the number of chimeric reads, as well as the number of TE-aligned reads and chimeric reads (Sup. Figure 14).

**Figure 7:**
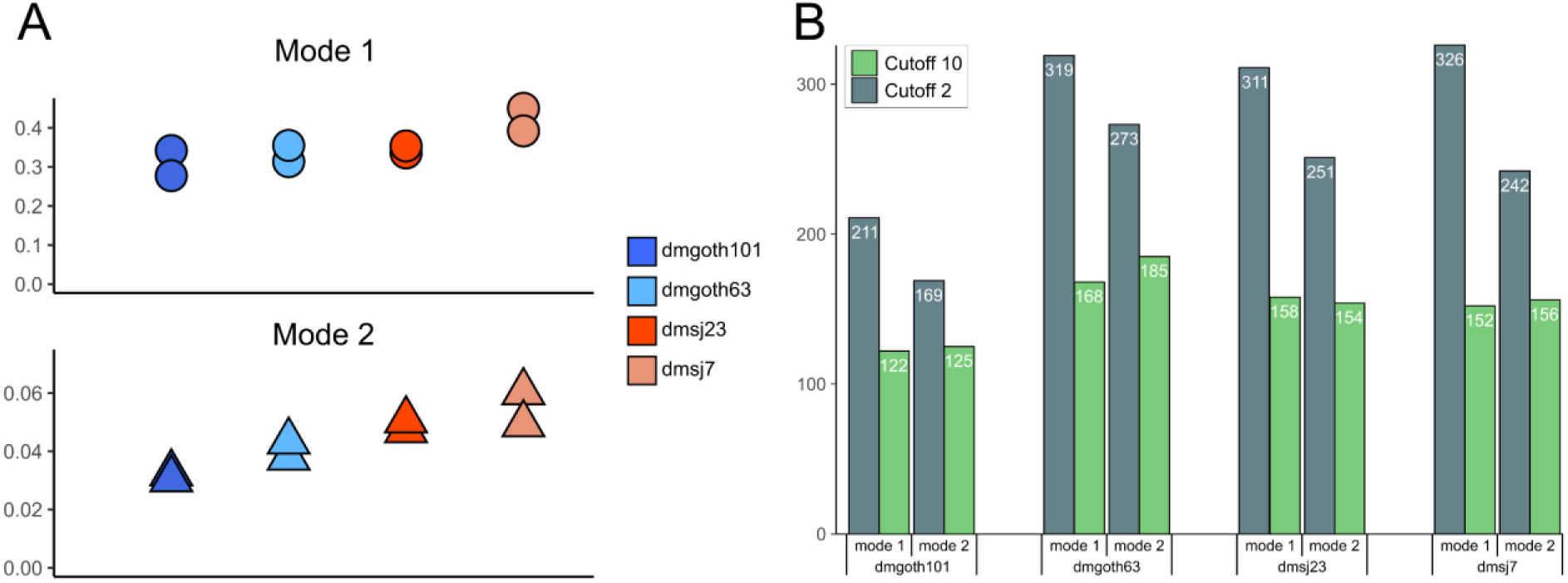
**A**) Total chimeric reads found in each RNA-seq replicate. **B**) Chimeric transcripts detected by both modes of ChimeraTE. The bars represent chimeras found in both RNA-seq replicates in the four *D. melanogaster* wild-type strains, applying two chimeric read cutoffs: 2 and 10.

To quantify ChimeraTE replicability between RNA-seq samples in both modes, as well as the impact of changes in the coverage thresholds, we performed a comparative analysis of chimeric transcripts using two thresholds of chimeric reads, 2 and 10. Overall, Mode 1 finds 291 and 150 chimeric transcripts per strain in both replicates (Figure 7B), using thresholds 2 and 10 respectively, whereas Mode 2 obtains 233 and 155. These results show that by increasing the chimeric read thresholds from 2 to 10, there is a decrease of 48.45% and 33.48% in the number of detected chimeric transcripts in Mode 1 and Mode 2. Therefore, even when found in both replicates, a substantial amount of chimeric transcripts is detected with low chimeric read coverage by both modes.

### Artifactual chimeric reads are largely reduced with RNA-seq replicability

In both Mode 1 and Mode 2, only chimeric transcripts found in both RNA-seq replicates were considered true chimeras. Chimeric transcripts found in only one replicate may exist due to pervasive transcription in one of the replicates, the lack of read coverage in one of the replicates to predict it, or they could be artifacts of cDNA library preparation, such as template switching and hybridization of templates followed by chimeric elongation (86–88). In addition, although comprising low rates with Illumina’s traditional bridge amplification, artifactual chimeric reads might be generated due to index hopping in multiplexed sequencing (37, 89). To quantify cases that may be artifacts, we aligned the RNA-seq from the four wild-type strains against their masked genomes to identify paired-end reads where each mate maps to genes from different chromosomes, or were aligned with an insert size > 500kb, and therefore are artifacts. It is important to highlight that these alignments do not differentiate artifactual reads produced during sequencing, and biological fusion gene products derived from the breakage and re-joining from different chromosomes, or chromosomal rearrangements (90, 91). In replicate 1 and replicate 2, we found ∼50,522 and ∼42,303 chimeric reads representing fusions of genes that are in different chromosomes, and ∼7,509, ∼5,334 farther away in the genome, with at least one chimeric read (Sup. Table 14). Such fusions were composed in average by ∼5,966 genes from different chromosomes and ∼3,234 genes farther away. However, we found only 498 (8.35%) of the artifactual chimeras from different chromosomes, and 212 (6.56%) genes farther away in both RNA-seq replicates with >= 2 chimeric reads (Sup. Table 14). In addition, these fusion genes generated by artifacts have high expression level (average FPKM = 722 per strain), reinforcing the hypothesis that high expressed genes are more likely to produce artifacts (86). Therefore, artifactual chimeric reads present in more than one replicate, and derived from genes and TEs, might have a lower prevalence, since the proportion of RNA-seq reads derived from TEs is very low (Figure 7A) in comparison to genes. Thus, in order to significantly reduce false positives, it is strongly recommended to use RNA-seq replicates, since artifactual chimeric reads exist in high frequency in highly expressed genes but only in one RNA-seq replicate. On the other hand, the proportion of the same chimeras found in more than one RNA-seq replicate with >= 2 chimeric reads is low.

### The high prevalence of *roo* elements in chimeric transcripts is due to ∼132 nt TE insertions embedded into exons

Several TE families have been associated with chimeric transcripts in *D. melanogaster* (31, 34). Here, we found 76 TE families producing at least one chimeric transcript in the four wild-type strains, representing 61.79% of all TE families annotated. We found a positive correlation between the frequency of TE families in chimeric transcripts and their respective abundance in the *D. melanogaster* genomes (*Pearson; r*=0.53; *p* < *2.2E-16*). In the TE-initiated transcripts, *S2*, *INE-1*, *hobo*, *1360* and *412* had the highest frequencies (Figure 8). In TE-exonized transcripts, the prevalence of *roo* elements is more pronounced in the embedded TEs group, comprising 50% of all chimeras, followed by *INE-1* elements, with 19%.

**Figure 8:**
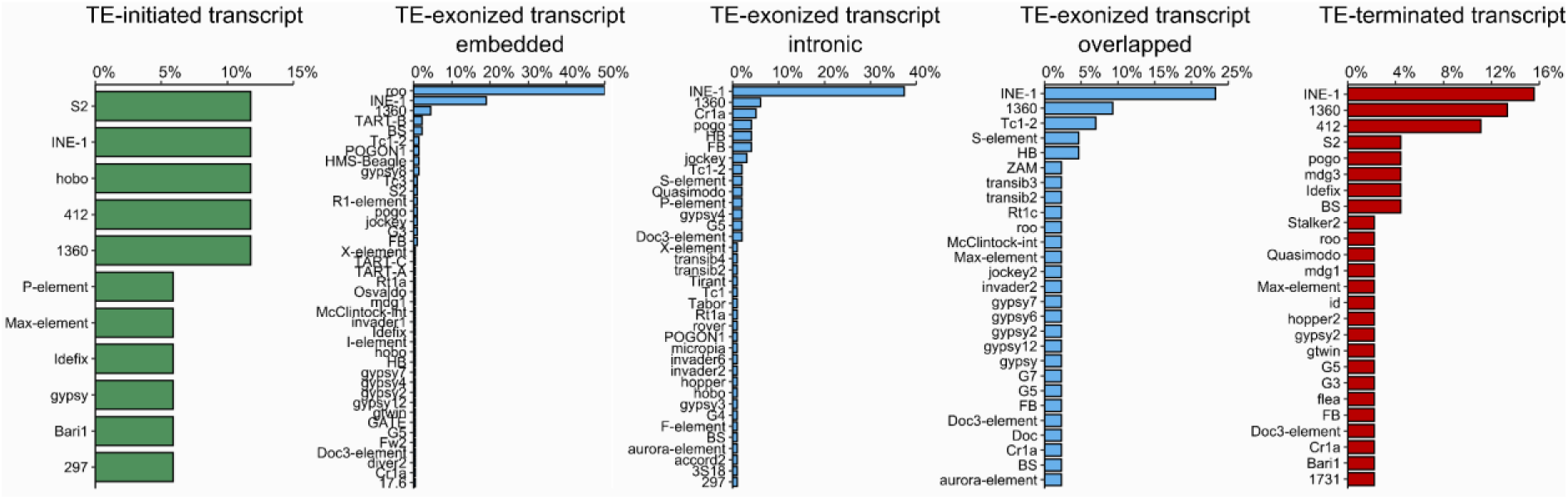
The frequency of the 76 TE families generating chimeric transcripts in the four wild-type strains. In chimeric transcripts derived from TEs near genes, *INE-1* was the most frequent (15%) in TE-terminated downstream, whereas for TE-initiated upstream, *S2, INE-1*, *hobo*, *1360* and *412* had the same frequency (11%). Regarding TEs inside genes, *the roo* element has the highest frequency of TE-exonized embedded, representing 50% of all chimeras. In TE-exonized intronic and overlapped, *INE-1* was the most frequent TE family.

Interestingly, for both TE-exonized intronic and overlapped, *INE-1* had the highest frequencies, with 37.37% and 23.25%, respectively (Figure 8); whereas *roo* element was observed in only one chimera (*Lk6* - *roo*, in dmgoth63 and dmsj23) as a TE-exonized overlapped. Notably, the *1360 P-element* is among the most frequent TE families in all TE-exonized categories. The other 38 families have an average frequency of 2%. Finally, in TE-terminated transcripts, *INE-1* is the most frequent TE, with 15.5% of all cases, followed by *1360* and *412*, with 13% and 11%, respectively. Taken together, these results suggest that *INE-1, S2, 412,* and *1360* are candidate TE families to investigate the exaptation of regulatory motifs since they were the most abundant in TE-initiated and TE-terminated transcripts. On the other hand, in TE-exonized transcripts, *roo* might have a potential for exaptation/domestication of both regulatory and protein domain motifs, since it is observed with high prevalence within exons (UTRs and CDSs). Regarding TE-exonized transcripts derived from TEs at intronic regions and overlapping with exons, *INE-1* and *1360* are the families with the highest frequency of intron retention, suggesting they might harbour splice sites allowing incorporation into the gene transcripts.

We then investigated whether exonized TE insertions generating TE-exonized transcripts could contribute to protein domains, especially *roo* due to its high prevalence in TE-exonized embedded chimeras. Through our analysis with *pfam-scan*, we found in total 28 genes with TE-exonized transcripts from TE insertions harbouring a TE protein domain, corresponding to an average of ∼16 per strain (Sup. Table 15). We assessed the completeness of these protein domains conserved on these insertions, assuming domains with > 75% of completeness as complete, and < 75% as partial. Our analysis revealed that ∼45% of all domains are likely to be conserved (Figure 9A), for which 92% correspond to catalytic domains, and 8% to zinc finger binding domains (PRE_C2HC – PF07530.14; zf-CCHC – PF00098.26). In addition, 17.69% of all TE-derived protein domains stem from embedded TE insertions, 27.43% from overlapped with exon, and 54.86% from intron regions. Furthermore, only one *roo* insertion had protein domains detected (Peptidase A17; DUF5641), which is embedded within CR46472 long noncoding RNA gene. Taking into account that 50% of the TE-exonized transcripts are derived from embedded *roo* insertions, this result suggests that despite their high frequency, they are old insertions without catalytic potential.

**Figure 9:**
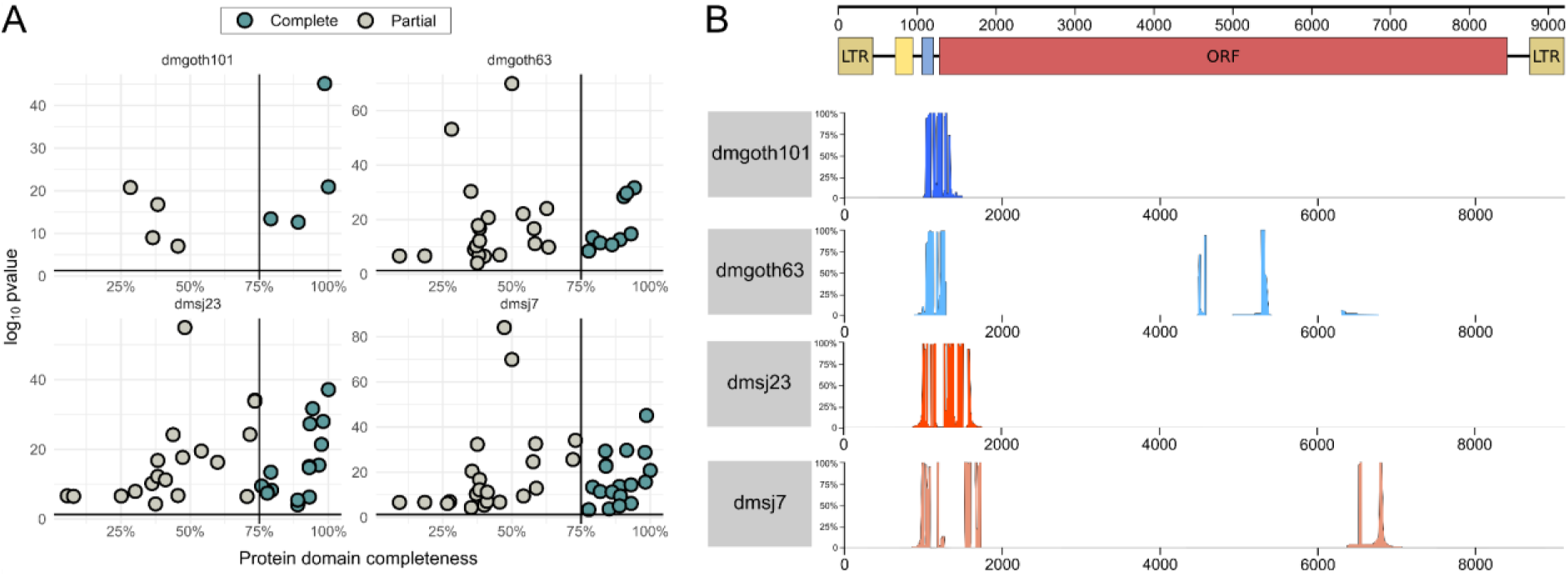
**A**) Completeness of protein domains identified in TE insertions generating TE-exonized transcripts, with p-value < 0.05. **B**) Alignment depth with *roo* consensus and embedded *roo* insertions generating TE-exonized transcripts. At the top, the scheme of the full-length *roo* element: brown boxes: LTRs; yellow box: first tandem repeats at 5’ UTR; blue box: second tandem repeat at 5’ UTR; red box: Open reading frame (ORF). The coverage depth of the multiple alignments between embedded *roo* insertions and the consensus is separated by strain.

Since we predict these chimeric transcripts only based on chimeric reads, we are not able to confirm whether these domains have been fully incorporated into gene transcripts, limiting our biological conclusions regarding catalytic or binding effects. There are only three chimeric transcripts with TE-derived protein domains for which we found the whole TE sequence incorporated into the transcript with ONT RNA-seq data (Sup. Figure 15). Among them, *CG4004* – *hobo*, and *nxf2* – *TART-A* are present into CDS regions, suggesting a putative functional role of the TE sequences. The third chimera, *CR42653* – *Doc*, is a non-coding pseudo-gene, which suggests that the TE-derived protein domain does not have a functional role.

TE insertions disrupting coding regions and promoters are more likely to be deleterious, being consequently eliminated by purifying selection (92), although several exaptation events have been documented (23, 93). The presence of *roo* elements in 50% of all TE-exonized embedded transcripts suggests a neutral or adaptive role of this family when incorporated into gene transcripts. *Roo* is an LTR retrotransposon, encoding three proteins: *gag*, *pol*, and *env*, which have been through domestication events from retroelements in many species, including *Drosophila* (8, 94, 95). It is the most abundant euchromatic TE family of *D. melanogaster* (1, 96, 97), and its insertions have been associated with modifications in the expression level of stress response genes due to the presence of TFBSs (98). They have also low enrichment of repressive histone marks (99, 100), potentially explaining their high transposition rates (101, 102). We analyzed the *roo* sequences that are embedded into exons of TE-exonized transcripts. The length of these chimeric insertions is ∼132 bp, confirming that they are old insertions since the full-length *roo* consensus sequence is 9,250 bp (Figure 9A). Subsequently, we analyzed whether these exonized *roo* insertions are donors of preferential motifs to the chimeric transcripts. All *roo* insertions stem from a specific region, between the 5’ UTR and the *roo* open-reading frame (Figure 9B). Compared to a *roo* consensus sequence, in dmgoth63 and dmsj7, most insertions show a deletion from the 5’ UTR up to the middle of the ORF. Despite the low nucleotide diversity of *roo* insertions in *the D. melanogaster* genome, the 5’ UTR region has a hypervariable region, including deletions and repeats, with several copies missing a tandem repeat of 99 bp (57, 103). It has been proposed that this region may have a role in *roo* transposition, by heterochromatinization, recruitment of RNA pol II, and interaction with other enzymes (103). Curiously, a study assessed the nucleotide diversity between *roo* insertions, and characterized the same region as a deletion hotspot (70). A recent work also observed this region as part of TE-exonized transcripts in a transcriptome-wide manner in *D. melanogaster* (73). Why this region has been maintained through evolution, and whether it could be adaptive or neutral, remains unclear.

### Polymorphic TE insertions in the wild-type strains generate 76 chimeric transcripts

Polymorphic TE insertions are common across *D. melanogaster* strains (78, 104). To quantify how many chimeric transcripts in the wild-type strains are derived from polymorphic TE insertions compared to the reference genome, we used the list of genes with TEs located 3 kb upstream, inside (introns and exons), and 3 kb downstream generated previously for the dm6 genome. These genes were selected as potential sources of chimeras and then compared with the list of chimeric transcripts generated by ChimeraTE Mode 1 in the four strains. We found that 76 genes (23.24%) with chimeric transcripts in the wild-derived strains were generated by TE insertions that are absent in the reference genome (Sup. Table 16). Twenty-seven TE families generate polymorphic chimeric transcripts: *roo* (44.15%), *412* (6.49%), *pogo* (6.49%), *POGON1* (6.49%), among others. Except for *INE-1*, for which we found two polymorphic chimeras, most of the other TE families are known to be active in *D. melanogaster* (57). *INE-1* chimeras are unexpected since it is an old and inactive TE family in *D. melanogaster* (105), however, *INE-1* polymorphism among *D. melanogaster* populations has previously been shown (106).

The study of TEs in wild-type strains offers new insights into their ability to provide genetic variability. Depending upon the position where TEs are inserted regarding genes, they can contribute to gene expression or protein sequence variation. We then compared whether polymorphic TE insertions generating chimeric transcripts are more likely inserted near or inside genes. We found that 5.19% correspond to TEs located upstream, 74.03% inside (intron/exons), and 20.78% downstream (Figure 10). In the genomic context, we observed ∼1,767 polymorphic TE insertions in these three regions regardless of the presence of chimeric transcripts, for which we found an equal distribution between TEs upstream (∼32%), TEs inside genes (∼35%), and TEs downstream (∼32%). Such result suggests that polymorphic insertions are not preferentially maintained by selection in specific gene locations, but chimeric transcripts from polymorphic TEs are more likely to be generated when they are within gene regions. In addition, we investigated whether the TE insertions absent from the reference genome could either be population-specific (Brazil or France) or strain-specific. The four TE-initiated transcripts derived from polymorphic TE insertions were strain-specific, two from dmgoth63 and two from dmsj7 (Figure 11A). For TEs inserted inside genes, 66.67% were found only in one strain. Only four chimeric transcripts are common to all strains *CG10543* - *roo*, *CG10077* - *roo*, *RASSF8* - *roo*, and *CG7239* - *roo* (Figure 11B). We did not find a TE-exonized derived from a TE insertion present only in the Brazilian strains, but we did find one for the French strains, the *simj – roo* chimera. Interestingly, nine chimeras were found between pairs of Brazilian and French strains (Figure 11B), indicating retention of ancestral polymorphisms in these populations due do incomplete lineage sorting (107). Finally, for TEs located downstream genes (Figure 11C), the chimera *Mps1–FB* is the only one found in both French strains, and absent from Brazilian strains, as we observed by Mode 2 (Figure 5). Taken together, these results reinforce the potential of TEs in generating genetic novelty between *D. melanogaster* strains.

**Figure 10:**
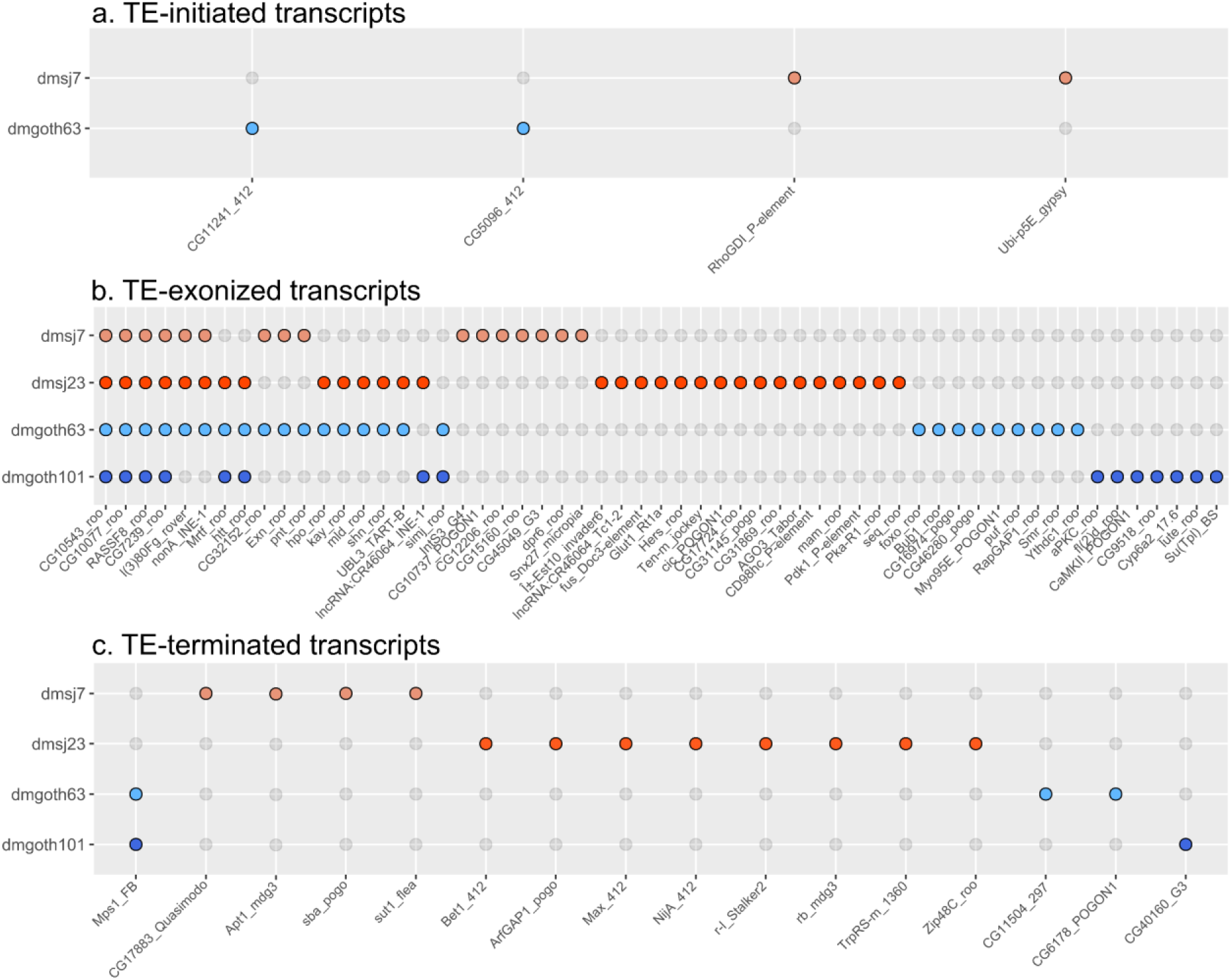
The 76 chimeric transcripts derived from TE insertions that are absent in the *dm6* reference genome, 5.19% of them correspond to TEs located upstream, 74.03% to TEs located inside genes (introns and exons), and 20.78% to TEs located downstream. **A**) TE upstream: Chimeric transcripts in which the TE is located up to 3kb upstream of the gene. **B**) TE inside: Chimeric transcripts with TE insertions located inside the gene region (exons and introns). There are four chimeric transcripts found in all strains, and one specific to French strains. **C**) TE downstream: Chimeric transcripts in which the TE is located up to 3kb downstream of the gene. Only *Mps1-FB* was specific to French strains, whereas all the other 15 chimeras are strain-specific.

**Figure 11:**
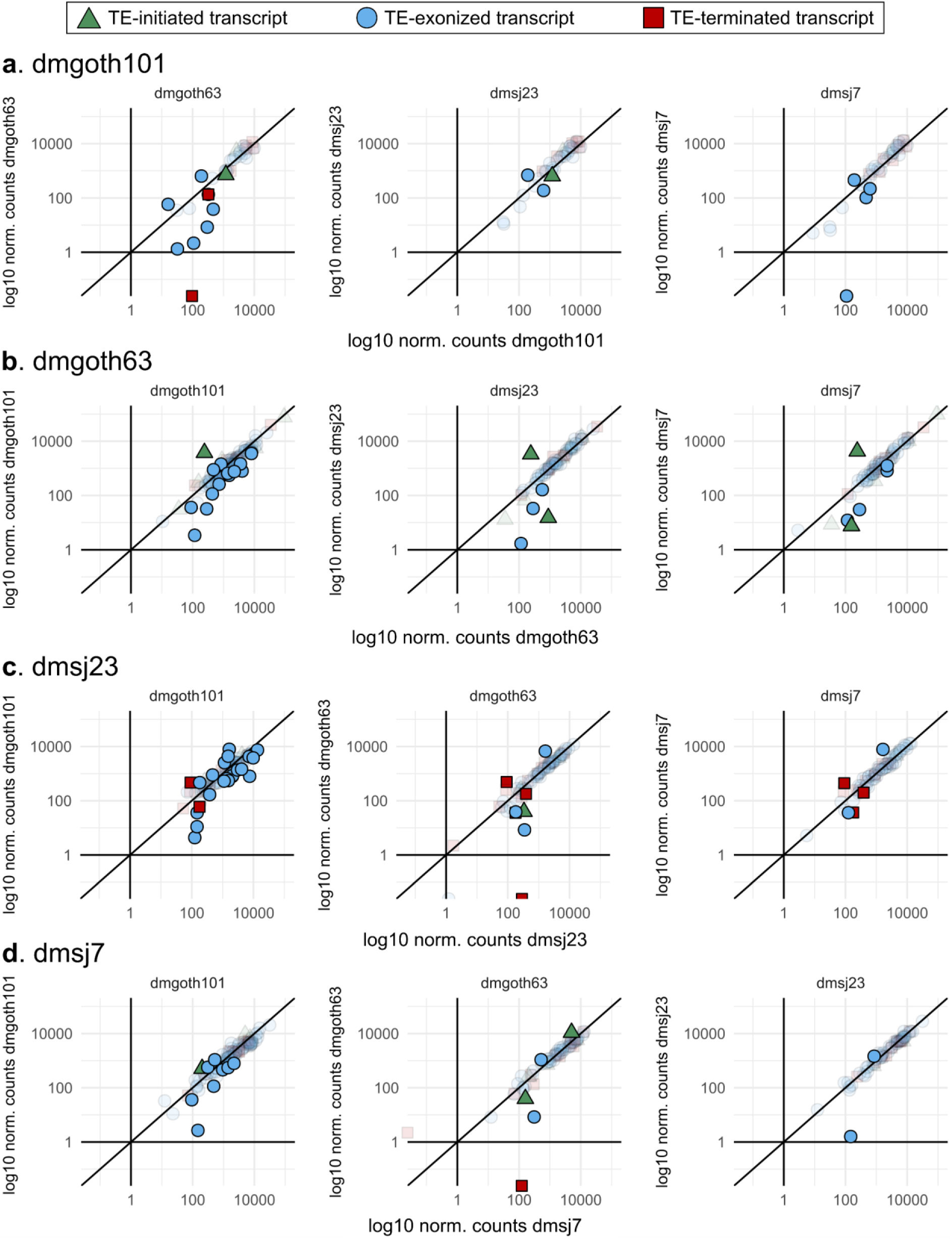
Normalized counts from DEseq2 indicating gene expression of genes generating polymorphic chimeric transcripts among the four wild-type strains. Colorful forms represent differentially expressed genes (adj. p-value < 0.05); transparent forms represent non-differentially expressed genes (adj. p-value). **A**) Expression level of genes producing chimeric transcripts only in dmgoth101, in comparison to dmgoth63, dmsj23, and dmsj7. **B**) Expression level of genes producing chimeric transcripts only in dmgoth63, in comparison to dmgoth101, dmsj23, and dmsj7. **C**) Expression level of genes producing chimeric transcripts only in dmsj23, in comparison to dmgoth101, dmgoth63, and dmsj7. **D**) Expression level of genes producing chimeric transcripts only in dmsj7, in comparison to dmgoth101, dmgoth63, and dmsj23. Overall, gene expression does not change when comparing strains with and without chimeric transcripts.

### Modification in gene expression is not a general rule for chimeric transcripts

TEs can modify gene expression when inserted inside and in their close vicinity. We hypothesized that genes with strain-specific chimeric transcripts may have differences in their expression levels in comparison to strains without the same chimeras. We first identified 67 strain-specific chimeras per strain, through pairwise comparisons. Then, through differential expression analysis between the four strains, we found that most genes have a similar expression level regardless of the presence of chimeric transcripts (Figure 11). In average, only 8 genes per strain were differentially expressed (Figure 11). Taking all these genes together, these chimeras are composed of 11.34% TE-initiated, 76.28% TE-exonized and 12.37% TE-terminated.

Among differentially expressed genes generating strain-specific chimeras, we observed that ∼47% had log2 fold-change < |2|, representing a mild transcription modulation. The other ∼53% (36 genes) had on average a log2 fold-change of ∼4.5 indicating higher expression when the chimera is present. Nevertheless, the gene expression quantification does not take into account differences between chimeric and non-chimeric isoforms. Therefore, we cannot differentiate gene expression differences due to chimeric isoform expression. We then focused on differentially expressed genes that are expressed in a given strain, but silent in the other (log2 normalized counts < 1). Only two chimeric cases fell into such category: TE-exonized CR45600 – *Tc1-2; gypsy7* in dmgoth101 compared to dmsj7 (Figure 11A); and the TE-terminated CG30428 – *INE-1* in dmgoth101, dmsj23, and dmsj7 compared to dmgoth63 (Figure 11A, C, D). To verify whether the chimeric transcript has a relatively high contribution to the gene expression, we compared the depth of uniquely mapped reads between exons and the TE insertion. For the CR45600 – *Tc1-2; gypsy7* chimera, we found a similar depth (Sup. Figure 16A), indicating that the chimeric transcript isoform represents a major contribution to the gene expression; whereas for CG30428 – *INE-1* chimera, the depth in *INE-1* is half of CG30428 exons (Sup. figure 16B), suggesting that the TE sequence contributes with only ∼50% of gene expression.

In dmgoth101, we found the gene *Cyp6a2* differentially expressed and involved in a TE-exonized transcript with a *17.6* insertion. Such chimera is absent from dmsj23, dmsj7, and dmgoth63 transcriptome. We checked whether *17.6* could be present on the genome of the other strains and we observed it only in dmgoth63, but undetected by ChimeraTE because *Cyp6a2* is not expressed. The presence of the *17.6* insertion within the *Cyp6a2* 3’UTR generating a chimeric transcript was initially proposed to cause the overexpression of this gene in flies resistant to xenobiotics (108), but later the opposite has been shown (109). Furthermore, regardless of *17.6* presence, the overexpression of this gene has been associated with xenobiotic resistance in *D. melanogaster* strains (110). Here, the differential expression analysis reveals that *Cyp6a2* is up-regulated in dmgoth101 in comparison to dmgoth63 (adj. p-value: 0.005; log2 FC: 4.38), but no significant differences were found when comparing with strains without the *17.6* insertion, dmsj23 (adj. p-value: 0.29), and dmsj7 (adj. pvalue: 0.15), reinforcing the lack of contribution of *17.6* insertion to *Cyp6a2* expression (109). Finally, the chimera derived from *Cyp6a14*, with a *1360* insertion embedded into the 3’ UTR has been found only in dmsj7, in comparison to the other three strains. This gene is associated with xenobiotic resistance (111), as *Cyp6a2*. Interestingly, our results reinforce previous findings showing that *Cyp* genes have accumulated more TE insertions when compared to a random sample of genes (112), potentially as a source of alternative regulatory motifs.

Collectively, the presence of chimeric transcripts might not drive gene differential expression. Despite eight differentially expressed genes per strain, the transcriptional activation of specific genes in one genotype may be determined by its genetic background, which can be associated with structural variants as TEs, but also includes pleiotropic effects from regulatory networks, cell physiology, and epigenetics (113). Nevertheless, given that TEs might provide TFBSs and epigenetic marks to chromatin accessibility, we propose these differentially expressed genes as candidates to investigate the domestication of regulatory networks in further studies, especially the ones with TEs inserted at regulatory regions.

### ChimeraTE uncovers chimeric transcripts in model and non-model species

We sought to demonstrate that both Modes of ChimeraTE are able to identify chimeric transcripts regardless of the species. Therefore, we used ChimeraTE with human, *A. thaliana* and *P. reticulata* (guppy fish) data. In Mode 1, the RNA-seq of the human K562 cell line revealed 67 TE-initiated, 14,999 TE-exonized, and 166 TE-terminated transcripts, derived from 5,170 genes (10.45% of total). Furthermore, Mode 2 revealed 5,632 chimeric transcripts. From these, 3,502 chimeras were detected from chimeric reads, 1,505 from transcriptome assembly, and 625 from double evidence. The overlap from chimeras found by both Modes was 2,503 (44.44%), suggesting a high amount of chimeric transcripts derived from TE insertions that are not present in the reference genome, or from TEs located farther away than the chosen 3 kb gene flanking regions. In *A. thaliana*, we used ChimeraTE with RNA-seq from leaf tissue. Our results from Mode 1 demonstrated that 266 genes (0.7% out of 38,319 generate 397 chimeras, corresponding to 24 TE-initiated, 338 TE-exonized, and 35 TE-terminated transcripts. Mode 2 has detected 141 genes generating chimeras, for which 63.12% were from chimeric reads, 24.11% from transcriptome assembly, and 12.77% from double evidence. Similar to human, Mode 2 found 39.72% of chimeras detected by Mode 1. Finally, the guppy transcriptome was analyzed with RNA-seq produced from ovary follicular tissue. For the first time, we identified 1,151 genes (5.85% of total genes) generating chimeric transcripts in this species. From these, 39 were TE-initiated, 1,507 TE-exonized, and 136 TE-terminated transcripts. Mode 2 revealed 640 genes with chimeras, having an overlap with 318 genes (49.69%) detected by Mode 1. Taking together our results from human, *A. thaliana*, guppy, and *D. melanogaster* data, we demonstrated that ChimeraTE can identify chimeras in different species.

We then analyzed how the genome size could affect ChimeraTE Mode 1 and Mode 2 in the processing time, and we have observed a positive correlation between them in Mode 1 (*Pearson; r = 0.99; p = 0.01*), but not in Mode 2 (Sup. figure 17). This is likely to be associated with the *de novo* transcriptome assembly applied in Mode 2, for which the processing time has a positive correlation with library size (48).

## CONCLUSION

In the last decades, RNA-seq has provided the opportunity to understand transcriptome plasticity, which can lead to phenotypic divergence, from related species or individuals from a population (114). Among several sources of modification in gene expression and novel isoform transcripts, TEs have been considered fundamental suppliers of transcriptome plasticity, participating in gene networks and incorporating either regulatory sequences or protein domains into gene transcripts. The identification of chimeric transcripts is an important step to the understanding of transcriptome plasticity, since they may be triggered by ectopic conditions, such as cancer, oxidative stress, and heat shock (26, 115, 116), which may lead to both detrimental and advantageous outcomes (117). Therefore, uncovering the extent of chimeric transcripts between individual/cell/strain transcriptomes is a crucial first step to investigate potential exaptation/domestication events, or gene disruption and loss of function.

Chimeric transcripts have been identified more recently by different methods exploiting RNA-seq data (34–36, 73), but none of them provided the possibility to predict chimeras from TEs that are absent from reference genomes. Here, we developed ChimeraTE, a pipeline able to identify chimeric transcripts from TEs that are absent from the reference genome. Mode 1 is a genome-guided approach and may be used either when the user is not interested in chimeras derived from TEs absent from the reference genome, or when the user has a high-quality genome assembly for each individual/cell/strain; whereas Mode 2 is a genome-blind approach, with the ability to predict chimeric transcripts without the assembled genome, but missing chimeras where the TE is smaller than the length of the reads.

We analyzed ovarian RNA-seq from four *D. melanogaster* wild-type strains, for which we have genome assemblies. Altogether, we found that ∼1.12% of all genes are generating chimeric transcripts in ovaries, following the proportion from previous studies obtained with ESTs, and RNA-seq in midbrain tissue (31, 34). Furthermore, our results revealed that 50% of all TE-exonized transcripts with TEs embedded derive from *roo* elements, and most specifically, a small region between tandem repeats in the 5’ UTR and the beginning of the *roo* ORF. These results suggest that these *roo* insertions have neutral or advantageous effects as they are maintained within these gene transcripts. However, we did not provide enough support to claim these *roo*-exonized transcripts as exaptation or domestication events, due to the lack of evidence regarding the functional role of these chimeras. Further studies must be performed to clarify this subject, mainly because chimeric transcripts detected by ChimeraTE can be degraded by surveillance pathways that degrade aberrant mRNA, such as non-sense-mediated mRNA decay (118), non-go decay (119), and non-stop decay (120).

Altogether, this new approach allows studying the impact of new mobilization events between populations or between treatment conditions, providing insights into biological questions from a broad community of researchers, ranging from cancer research, to population transcriptomics, and adaptation studies. ChimeraTE implementation will be useful for the next discoveries regarding the evolutionary role of TEs and their impact on the host transcriptome.

## AVAILABILITY

ChimeraTE is an open-source collaborative initiative available in the GitHub repository (https://github.com/OliveiraDS-hub/ChimeraTE).

## ACCESSION NUMBERS

The RNA-seq data used in this study are available in the NCBI BioProject database (https://www.ncbi.nlm.nih.gov/bioproject/), under PRJNA795668 accession, while the long-read ONT data is available under PRJNA956863.

## ACKNOWLEDGEMENT

We thank Josefa González, Marta Coronado-Zamora, Dixie Mager, and Vincent Lacroix for useful discussions and advice. This work was performed using the computing facilities of the CC LBBE/PRABI.

## REFERENCES

1. Quesneville, H., Bergman, C.M., Andrieu, O., Autard, D., Nouaud, D., Ashburner, M. and Anxolabehere, D. (2005) Combined Evidence Annotation of Transposable Elements in Genome Sequences. PLoS Comp Biol, 1, e22.

2. International Human Genome Sequencing Consortium, Whitehead Institute for Biomedical Research, Center for Genome Research:, Lander, E.S., Linton, L.M., Birren, B., Nusbaum, C., Zody, M.C., Baldwin, J., Devon, K., Dewar, K., et al. (2001) Initial sequencing and analysis of the human genome. Nature, 409, 860–921.

3. Schnable, P.S., Ware, D., Fulton, R.S., Stein, J.C., Wei, F., Pasternak, S., Liang, C., Zhang, J., Fulton, L., Graves, T.A., et al. (2009) The B73 Maize Genome: Complexity, Diversity, and Dynamics. Science, 326, 1112–1115.

4. Sotero-Caio, C.G., Platt, R.N., Suh, A. and Ray, D.A. (2017) Evolution and Diversity of Transposable Elements in Vertebrate Genomes. Genome Biology and Evolution, 9, 161–177.

5. Danilevskaya, O.N., Arkhipova, I.R., Pardue, M.L. and Traverse, K.L. (1997) Promoting in Tandem: The Promoter for Telomere Transposon HeT-A and Implications for the Evolution of Retroviral LTRs. Cell, 88, 647–655.

6. Kapitonov, V.V. and Jurka, J. (2005) RAG1 Core and V(D)J Recombination Signal Sequences Were Derived from Transib Transposons. PLoS Biol, 3, e181.

7. Kapitonov, V.V. and Koonin, E.V. (2015) Evolution of the RAG1-RAG2 locus: both proteins came from the same transposon. Biol Direct, 10, 20.

8. Volff, J.-N. (2006) Turning junk into gold: domestication of transposable elements and the creation of new genes in eukaryotes. Bioessays, 28, 913–922.

9. Babaian, A., Romanish, M.T., Gagnier, L., Kuo, L.Y., Karimi, M.M., Steidl, C. and Mager, D.L. (2016) Onco-exaptation of an endogenous retroviral LTR drives IRF5 expression in Hodgkin lymphoma. Oncogene, 35, 2542–2546.

10. Daborn, P.J., Yen, J.L., Bogwitz, M.R., Le Goff, G., Feil, E., Jeffers, S., Tijet, N., Perry, T., Heckel, D., Batterham, P., et al. (2002) A Single P450 Allele Associated with Insecticide Resistance in *Drosophila*. Science, 297, 2253–2256.

11. Jordan, I.K., Rogozin, I.B., Glazko, G.V. and Koonin, E.V. (2003) Origin of a substantial fraction of human regulatory sequences from transposable elements. Trends in Genetics, 19, 68–72.

12. Mateo, L., Ullastres, A. and González, J. (2014) A Transposable Element Insertion Confers Xenobiotic Resistance in Drosophila. PLoS Genet, 10, e1004560.

13. Modzelewski, A.J., Shao, W., Chen, J., Lee, A., Qi, X., Noon, M., Tjokro, K., Sales, G., Biton, A., Anand, A., et al. (2021) A mouse-specific retrotransposon drives a conserved Cdk2ap1 isoform essential for development. Cell, 184, 5541-5558.e22.

14. Capy, P. (2021) Taming, Domestication and Exaptation: Trajectories of Transposable Elements in Genomes. Cells, 10, 3590.

15. Fueyo, R., Judd, J., Feschotte, C. and Wysocka, J. (2022) Roles of transposable elements in the regulation of mammalian transcription. Nat Rev Mol Cell Biol, 10.1038/s41580-022-00457-y.

16. Lamprecht, B., Walter, K., Kreher, S., Kumar, R., Hummel, M., Lenze, D., Köchert, K., Bouhlel, M.A., Richter, J., Soler, E., et al. (2010) Derepression of an endogenous long terminal repeat activates the CSF1R proto-oncogene in human lymphoma. Nat Med, 16, 571–579.

17. McGinnis, W., Shermoen, A.W. and Beckendorf, S.K. (1983) A transposable element inserted just 5′ to a Drosophila glue protein gene alters gene expression and chromatin structure. Cell, 34, 75–84.

18. Almeida, L.M., Amaral, M.E.J., Silva, I.T., Silva Jr, W.A., Riggs, P.K. and Carareto, C.M. (2008) Report of a chimeric origin of transposable elements in a bovine-coding gene. Genet. Mol. Res., 7, 107–116.

19. Sela, N., Mersch, B., Hotz-Wagenblatt, A. and Ast, G. (2010) Characteristics of Transposable Element Exonization within Human and Mouse. PLoS ONE, 5, e10907.

20. Sorek, R. (2007) The birth of new exons: Mechanisms and evolutionary consequences. RNA, 13, 1603–1608.

21. Bogwitz, M.R., Chung, H., Magoc, L., Rigby, S., Wong, W., O’Keefe, M., McKenzie, J.A., Batterham, P. and Daborn, P.J. (2005) Cyp12a4 confers lufenuron resistance in a natural population of Drosophila melanogaster. Proceedings of the National Academy of Sciences, 102, 12807–12812.

22. Farré, D., Engel, P. and Angulo, A. (2016) Novel Role of 3’UTR-Embedded Alu Elements as Facilitators of Processed Pseudogene Genesis and Host Gene Capture by Viral Genomes. PLoS ONE, 11, e0169196.

23. Magwire, M.M., Bayer, F., Webster, C.L., Cao, C. and Jiggins, F.M. (2011) Successive Increases in the Resistance of Drosophila to Viral Infection through a Transposon Insertion Followed by a Duplication. PLoS Genet, 7, e1002337.

24. Cordaux, R., Udit, S., Batzer, M.A. and Feschotte, C. (2006) Birth of a chimeric primate gene by capture of the transposase gene from a mobile element. Proc. Natl. Acad. Sci. U.S.A., 103, 8101–8106.

25. Ehrlich, M. (2009) DNA hypomethylation in cancer cells. Epigenomics, 1, 239–259.

26. Babaian, A. and Mager, D.L. (2016) Endogenous retroviral promoter exaptation in human cancer. Mobile DNA, 7, 24.

27. Lock, F.E., Rebollo, R., Miceli-Royer, K., Gagnier, L., Kuah, S., Babaian, A., Sistiaga-Poveda, M., Lai, C.B., Nemirovsky, O., Serrano, I., et al. (2014) Distinct isoform of FABP7 revealed by screening for retroelement-activated genes in diffuse large B-cell lymphoma. Proceedings of the National Academy of Sciences, 111, E3534–E3543.

28. Shah, N.M., Jang, H.J., Liang, Y., Maeng, J.H., Tzeng, S.-C., Wu, A., Basri, N.L., Qu, X., Fan, C., Li, A., et al. (2023) Pan-cancer analysis identifies tumor-specific antigens derived from transposable elements. Nat Genet, 55, 631–639.

29. Faulkner, G.J., Kimura, Y., Daub, C.O., Wani, S., Plessy, C., Irvine, K.M., Schroder, K., Cloonan, N., Steptoe, A.L., Lassmann, T., et al. (2009) The regulated retrotransposon transcriptome of mammalian cells. Nat Genet, 41, 563–571.

30. Babarinde, I.A., Ma, G., Li, Y., Deng, B., Luo, Z., Liu, H., Abdul, M.M., Ward, C., Chen, M., Fu, X., et al. (2021) Transposable element sequence fragments incorporated into coding and noncoding transcripts modulate the transcriptome of human pluripotent stem cells. Nucleic Acids Research, 49, 9132–9153.

31. Lipatov, M., Lenkov, K., Petrov, D.A. and Bergman, C.M. (2005) Paucity of chimeric gene-transposable element transcripts in the Drosophila melanogaster genome. BMC Biol, 3, 24.

32. Batut, P. and Gingeras, T.R. (2013) RAMPAGE: Promoter Activity Profiling by Paired-End Sequencing of 5′-Complete cDNAs. Current Protocols in Molecular Biology, 104.

33. Batut, P., Dobin, A., Plessy, C., Carninci, P. and Gingeras, T.R. (2013) High-fidelity promoter profiling reveals widespread alternative promoter usage and transposon-driven developmental gene expression. Genome Research, 23, 169–180.

34. Treiber, C.D. and Waddell, S. (2020) Transposon expression in the *Drosophila* brain is driven by neighboring genes and diversifies the neural transcriptome. Genome Res., 30, 1559–1569.

35. Pinson, M.-E., Pogorelcnik, R., Court, F., Arnaud, P. and Vaurs-Barrière, C. (2018) CLIFinder: identification of LINE-1 chimeric transcripts in RNA-seq data. Bioinformatics, 34, 688–690.

36. Babaian, A., Thompson, I.R., Lever, J. and Gagnier, L. (2019) LIONS: Analysis Suite for Detecting and Quan-tifying Transposable Element Initiated Tran-scription from RNA-seq. Bioinformatics, 35, 3839–3841.

37. Kircher, M., Sawyer, S. and Meyer, M. (2012) Double indexing overcomes inaccuracies in multiplex sequencing on the Illumina platform. Nucleic Acids Research, 40, e3–e3.

38. Evrony, G.D., Lee, E., Park, P.J. and Walsh, C.A. (2016) Resolving rates of mutation in the brain using single-neuron genomics. eLife, 5, e12966.

39. Quail, M.A., Kozarewa, I., Smith, F., Scally, A., Stephens, P.J., Durbin, R., Swerdlow, H. and Turner, D.J. (2008) A large genome center’s improvements to the Illumina sequencing system. Nat Methods, 5, 1005–1010.

40. Treiber, C.D. and Waddell, S. (2017) Resolving the prevalence of somatic transposition in Drosophila. eLife, 6, e28297.

41. Martin Cerezo, M.L., Raval, R., de Haro Reyes, B., Kucka, M., Chan, F.Y. and Bryk, J. (2022) Identification and quantification of chimeric sequencing reads in a highly multiplexed RAD -seq protocol. Molecular Ecology Resources, 10.1111/1755-0998.13661.

42. Trapnell, C., Roberts, A., Goff, L., Pertea, G., Kim, D., Kelley, D.R., Pimentel, H., Salzberg, S.L., Rinn, J.L. and Pachter, L. (2012) Differential gene and transcript expression analysis of RNA-seq experiments with TopHat and Cufflinks. Nat Protoc, 7, 562–578.

43. Li, H., Handsaker, B., Wysoker, A., Fennell, T., Ruan, J., Homer, N., Marth, G., Abecasis, G., Durbin, R., and 1000 Genome Project Data Processing Subgroup (2009) The Sequence Alignment/Map format and SAMtools. Bioinformatics, 25, 2078–2079.

44. Quinlan, A.R. and Hall, I.M. (2010) BEDTools: a flexible suite of utilities for comparing genomic features. Bioinformatics, 26, 841–842.

45. Dale, R.K., Pedersen, B.S. and Quinlan, A.R. (2011) Pybedtools: a flexible Python library for manipulating genomic datasets and annotations. Bioinformatics, 27, 3423–3424.

46. Langmead, B. and Salzberg, S.L. (2012) Fast gapped-read alignment with Bowtie 2. Nat Methods, 9, 357–359.

47. Roberts, A. and Pachter, L. (2013) Streaming fragment assignment for real-time analysis of sequencing experiments. Nat Methods, 10, 71–73.

48. Grabherr, M.G., Haas, B.J., Yassour, M., Levin, J.Z., Thompson, D.A., Amit, I., Adiconis, X., Fan, L., Raychowdhury, R., Zeng, Q., et al. (2011) Full-length transcriptome assembly from RNA-Seq data without a reference genome. Nat Biotechnol, 29, 644–652.

49. SMIT, Arian FA (2004) RepeatMasker Open 3.0.

50. Storer, J., Hubley, R., Rosen, J., Wheeler, T.J. and Smit, A.F. (2021) The Dfam community resource of transposable element families, sequence models, and genome annotations. Mobile DNA, 12, 2.

51. Altschul, S. (1997) Gapped BLAST and PSI-BLAST: a new generation of protein database search programs. Nucleic Acids Research, 25, 3389–3402.

52. Fablet, M., Salcez-Ortiz, J., Jacquet, A., Menezes, B.F., Dechaud, C., Veber, P., Noûs, C., Rebollo, R. and Vieira, C. (2022) A quantitative, genome-wide analysis in Drosophila reveals transposable elements’ influence on gene expression is species-specific. bioRxiv.

53. Bolger, A.M., Lohse, M. and Usadel, B. (2014) Trimmomatic: a flexible trimmer for Illumina sequence data. Bioinformatics, 30, 2114–2120.

54. Mohamed, M., Dang, N.T.-M., Ogyama, Y., Burlet, N., Mugat, B., Boulesteix, M., Mérel, V., Veber, P., Salces-Ortiz, J., Severac, D., et al. (2020) A Transposon Story: From TE Content to TE Dynamic Invasion of Drosophila Genomes Using the Single-Molecule Sequencing Technology from Oxford Nanopore. Cells, 9, 1776.

55. Shumate, A. and Salzberg, S.L. (2021) Liftoff: accurate mapping of gene annotations. Bioinformatics, 37, 1639–1643.

56. Wicker, T., Sabot, F., Hua-Van, A., Bennetzen, J.L., Capy, P., Chalhoub, B., Flavell, A., Leroy, P., Morgante, M., Panaud, O., et al. (2007) A unified classification system for eukaryotic transposable elements. Nat Rev Genet, 8, 973–982.

57. Lerat, E., Rizzon, C. and Biémont, C. (2003) Sequence Divergence Within Transposable Element Families in the *Drosophila melanogaster* Genome. Genome Res., 13, 1889–1896.

58. Flutre, T., Duprat, E., Feuillet, C. and Quesneville, H. (2011) Considering Transposable Element Diversification in De Novo Annotation Approaches. PLoS ONE, 6, e16526.

59. Flutre, T., Permal, E. and Quesneville, H. (2012) Transposable Element Annotation in Completely Sequenced Eukaryote Genomes. In Grandbastien, M.-A., Casacuberta, J.M. (eds), Plant Transposable Elements, Topics in Current Genetics. Springer Berlin Heidelberg, Berlin, Heidelberg, Vol. 24, pp. 17–39.

60. Benson, G. (1999) Tandem repeats finder: a program to analyze DNA sequences. Nucleic Acids Research, 27, 573–580.

61. dos Santos, G., Schroeder, A.J., Goodman, J.L., Strelets, V.B., Crosby, M.A., Thurmond, J., Emmert, D.B., Gelbart, W.M., and the FlyBase Consortium (2015) FlyBase: introduction of the Drosophila melanogaster Release 6 reference genome assembly and large-scale migration of genome annotations. Nucleic Acids Research, 43, D690–D697.

62. Lee, C.M., Barber, G.P., Casper, J., Clawson, H., Diekhans, M., Gonzalez, J.N., Hinrichs, A.S., Lee, B.T., Nassar, L.R., Powell, C.C., et al. (2019) UCSC Genome Browser enters 20th year. Nucleic Acids Research, 10.1093/nar/gkz1012.

63. Thorvaldsdottir, H., Robinson, J.T. and Mesirov, J.P. (2013) Integrative Genomics Viewer (IGV): high-performance genomics data visualization and exploration. Briefings in Bioinformatics, 14, 178–192.

64. Rice, P., Longden, I. and Bleasby, A. EMBOSS: The European Molecular Biology Open Software Suite. Trends in Genetics, 16, 276–277.

65. Mistry, J., Chuguransky, S., Williams, L., Qureshi, M., Salazar, G.A., Sonnhammer, E.L.L., Tosatto, S.C.E., Paladin, L., Raj, S., Richardson, L.J., et al. (2021) Pfam: The protein families database in 2021. Nucleic Acids Research, 49, D412–D419.

66. Paysan-Lafosse, T., Blum, M., Chuguransky, S., Grego, T., Pinto, B.L., Salazar, G.A., Bileschi, M.L., Bork, P., Bridge, A., Colwell, L., et al. (2023) InterPro in 2022. Nucleic Acids Research, 51, D418–D427.

67. Edgar, R.C. (2004) MUSCLE: multiple sequence alignment with high accuracy and high throughput. Nucleic Acids Research, 32, 1792–1797.

68. Zea, D.J., Anfossi, D., Nielsen, M. and Marino-Buslje, C. (2017) MIToS.jl: mutual information tools for protein sequence analysis in the Julia language. Bioinformatics, 33, 564–565.

69. Love, M.I., Huber, W. and Anders, S. (2014) Moderated estimation of fold change and dispersion for RNA-seq data with DESeq2. Genome Biol, 15, 550.

70. Kaminker, J.S., Bergman, C.M., Kronmiller, B., Svirskas, R., Patel, S., Frise, E., Lewis, S.E., Rubin, G.M., Ashburner, M. and Celniker, S.E. (2002) The transposable elements of the Drosophila melanogaster euchromatin: a genomics perspective. Genome Biology, 3, 1–20.

71. Gatti, M. and Pimpinelli, S. (1992) Functional elements in Drosophila melanogaster heterochromatin. Annual review of genetics, 26, 239–276.

72. Carmena, M. and Gonzfilez, C. (1995) Transposable elements map in a conserved pattern of distribution extending from beta-heterochromatin to centromeres in Drosophilamelanogaster. Chromosoma, 103, 676–684.

73. Coronado-Zamora, M. and González, J. (2022) Transposons contribute to the functional diversification of the head, gut, and ovary transcriptomes across *Drosophila* natural strains. bioRxiv, 10.1101/2022.12.02.518890.

74. Bailly-Bechet, M., Haudry, A. and Lerat, E. (2014) “One code to find them all”: a perl tool to conveniently parse RepeatMasker output files. Mobile DNA, 5, 13.

75. Kapun, M., Barrón, M.G., Staubach, F., Obbard, D.J., Wiberg, R.A.W., Vieira, J., Goubert, C., Rota-Stabelli, O., Kankare, M., Bogaerts-Márquez, M., et al. (2020) Genomic Analysis of European Drosophila melanogaster Populations Reveals Longitudinal Structure, Continent-Wide Selection, and Previously Unknown DNA Viruses. Molecular Biology and Evolution, 37, 2661–2678.

76. Rech, G.E., Bogaerts-Márquez, M., Barrón, M.G., Merenciano, M., Villanueva-Cañas, J.L., Horváth, V., Fiston-Lavier, A.-S., Luyten, I., Venkataram, S., Quesneville, H., et al. (2019) Stress response, behavior, and development are shaped by transposable element-induced mutations in Drosophila. PLoS Genet, 15, e1007900.

77. Rech, G.E., Radío, S., Guirao-Rico, S., Aguilera, L., Horvath, V., Green, L., Lindstadt, H., Jamilloux, V., Quesneville, H. and González, J. (2022) Population-scale long-read sequencing uncovers transposable elements associated with gene expression variation and adaptive signatures in Drosophila. Nat Commun, 13, 1948.

78. Vieira, C., Lepetit, D., Dumont, S. and Biemont, C. (1999) Wake up of transposable elements following Drosophila simulans worldwide colonization. Molecular Biology and Evolution, 16, 1251–1255.

79. Lima, L., Sinaimeri, B., Sacomoto, G., Lopez-Maestre, H., Marchet, C., Miele, V., Sagot, M.-F. and Lacroix, V. (2017) Playing hide and seek with repeats in local and global de novo transcriptome assembly of short RNA-seq reads. Algorithms Mol Biol, 12, 2.

80. Lu, Z., Marand, A.P., Ricci, W.A., Ethridge, C.L., Zhang, X. and Schmitz, R.J. (2019) The prevalence, evolution and chromatin signatures of plant regulatory elements. Nat. Plants, 5, 1250–1259.

81. Bakoulis, S., Krautz, R., Alcaraz, N., Salvatore, M. and Andersson, R. (2022) Endogenous retroviruses co-opted as divergently transcribed regulatory elements shape the regulatory landscape of embryonic stem cells. Nucleic Acids Research, 50, 2111– 2127.

82. Lahens, N.F., Kavakli, I.H., Zhang, R., Hayer, K., Black, M.B., Dueck, H., Pizarro, A., Kim, J., Irizarry, R., Thomas, R.S., et al. (2014) IVT-seq reveals extreme bias in RNA sequencing. Genome Biology, 15, 15.

83. Wang, S. and Gribskov, M. (2016) Comprehensive evaluation of *de novo* transcriptome assembly programs and their effects on differential gene expression analysis. Bioinformatics, 33, 327–333.

84. Rebollo, R., Cumunel, E., Mary, A., Burlet, N., Gillet, B., Hughes, S., Oliveira, D.S., Goubert, C., Fablet, M., Vieira, C., et al. (2023) Detection and identification of transposable element transcripts using Long Read RNA-seq in Drosophila germline tissues. bioRxiv.

85. Ura, H., Togi, S. and Niida, Y. (2022) A comparison of mRNA sequencing (RNA-Seq) library preparation methods for transcriptome analysis. BMC Genomics, 23, 303.

86. Annala, M.J., Parker, B.C., Zhang, W. and Nykter, M. (2013) Fusion genes and their discovery using high throughput sequencing. Cancer Letters, 340, 192–200.

87. Kanagawa, T. (2003) Bias and artifacts in multitemplate polymerase chain reactions (PCR). Journal of bioscience and bioengineering, 96, 317–323.

88. Houseley, J. and Tollervey, D. (2010) Apparent Non-Canonical Trans-Splicing Is Generated by Reverse Transcriptase In Vitro. PLoS ONE, 5, e12271.

89. Van Der Valk, T., Vezzi, F., Ormestad, M., Dalén, L. and Guschanski, K. (2020) Index hopping on the Illumina HiseqX platform and its consequences for ancient DNA studies. Mol Ecol Resour, 20, 1171–1181.

90. Mitelman, F. (2012) Mitelman database of chromosome aberrations and gene fusions in cancer. http://cgap.nci.nih.gov/Chromosomes/Mitelman.

91. Zhou, Y., Zhang, C., Zhang, L., Ye, Q., Liu, N., Wang, M., Long, G., Fan, W., Long, M. and Wing, R.A. (2022) Gene fusion as an important mechanism to generate new genes in the genus Oryza. Genome Biol, 23, 130.

92. Cridland, J.M., Macdonald, S.J., Long, A.D. and Thornton, K.R. (2013) Abundance and Distribution of Transposable Elements in Two Drosophila QTL Mapping Resources. Molecular Biology and Evolution, 30, 2311–2327.

93. Lerman, D.N. and Feder, M.E. (2005) Naturally Occurring Transposable Elements Disrupt hsp70 Promoter Function in Drosophila melanogaster. Molecular Biology and Evolution, 22, 776–783.

94. Nefedova, L.N., Kuzmin, I.V., Makhnovskii, P.A. and Kim, A.I. (2014) Domesticated retroviral GAG gene in Drosophila: New functions for an old gene. Virology, 450–451, 196–204.

95. Malik, H.S. and Henikoff, S. (2005) Positive Selection of Iris, a Retroviral Envelope– Derived Host Gene in Drosophila melanogaster. PLoS Genet, 1, e44.

96. Rahman, R., Chirn, G., Kanodia, A., Sytnikova, Y.A., Bergman, C.M. and Lau, N.C. (2015) Unique transposon landscapes are pervasive across Drosophila melanogaster genomes. Nucleic Acids Research, 43, 10655–10672.

97. Vieira, C. and Biemont, C. (2004) Transposable element dynamics in two sibling species: Drosophila melanogaster and Drosophila simulans. Genetica, 120, 115–123.

98. Merenciano, M., Ullastres, A., de Cara, M.A.R., Barrón, M.G. and González, J. (2016) Multiple Independent Retroelement Insertions in the Promoter of a Stress Response Gene Have Variable Molecular and Functional Effects in Drosophila. PLoS Genet, 12, e1006249.

99. Rebollo, R., Horard, B., Begeot, F., Delattre, M., Gilson, E. and Vieira, C. (2012) A Snapshot of Histone Modifications within Transposable Elements in Drosophila Wild Type Strains. PLoS ONE, 7, e44253.

100. Yasuhara, J.C. and Wakimoto, B.T. (2008) Molecular Landscape of Modified Histones in Drosophila Heterochromatic Genes and Euchromatin-Heterochromatin Transition Zones. PLoS Genet, 4, e16.

101. Díaz-González, J., Vázquez, J.F., Albornoz, J. and Domínguez, A. (2011) Long-term evolution of the *roo* transposable element copy number in mutation accumulation lines of *Drosophila melanogaster*. Genet. Res., 93, 181–187.

102. Díaz-González, J., Domínguez, A. and Albornoz, J. (2010) Genomic distribution of retrotransposons 297, 1731, copia, mdg1 and roo in the Drosophila melanogaster species subgroup. Genetica, 138, 579–586.

103. Díaz-González, J. and Domínguez, A. (2020) Different structural variants of roo retrotransposon are active in Drosophila melanogaster. Gene, 741, 144546.

104. Barrón, M.G., Fiston-Lavier, A.-S., Petrov, D.A. and González, J. (2014) Population Genomics of Transposable Elements in *Drosophila*. Annu. Rev. Genet., 48, 561–581.

105. Kapitonov, V.V. and Jurka, J. (2003) Molecular paleontology of transposable elements in the *Drosophila melanogaster* genome. Proc. Natl. Acad. Sci. U.S.A., 100, 6569– 6574.

106. Kofler, R., Betancourt, A.J. and Schlötterer, C. (2012) Sequencing of Pooled DNA Samples (Pool-Seq) Uncovers Complex Dynamics of Transposable Element Insertions in Drosophila melanogaster. PLoS Genet, 8, e1002487.

107. Maddison, W.P. and Knowles, L.L. (2006) Inferring Phylogeny Despite Incomplete Lineage Sorting. Systematic Biology, 55, 21–30.

108. Waters, L.C. (1992) Possible involvement of the long terminal repeat of transposable element 17.6 in regulating expression of an insecticide resistance-associated P450 gene in Drosophila. Proc. Natl. Acad. Sci. USA, 89, 4855–4859.

109. Delpuech, J.M., Aquadro, C.F. and Roush, R.T. (1993) Noninvolvement of the long terminal repeat of transposable element 17.6 in insecticide resistance in Drosophila. Proc. Natl. Acad. Sci. U.S.A., 90, 5643–5647.

110. Brun, A., Cuany, A., Le Mouel, T., Berge, J. and Amichot, M. (1996) Inducibility of the Drosophila melanogaster cytochrome P450 gene, CYP6A2, by phenobarbital in insecticide susceptible or resistant strains. Insect Biochemistry and Molecular Biology, 26, 697–703.

111. Seong, K.M., Coates, B.S., Berenbaum, M.R., Clark, J.M. and Pittendrigh, B.R. (2018) Comparative CYP-omic analysis between the DDT-susceptible and -resistant *DROSOPHILA MELANOGASTER* strains *91-C* and *91-R*. Pest. Manag. Sci., 74, 2530– 2543.

112. Carareto, C.M.A., Hernandez, E.H. and Vieira, C. (2014) Genomic regions harboring insecticide resistance-associated Cyp genes are enriched by transposable element fragments carrying putative transcription factor binding sites in two sibling Drosophila species. Gene, 537, 93–99.

113. Hill, M.S., Vande Zande, P. and Wittkopp, P.J. (2021) Molecular and evolutionary processes generating variation in gene expression. Nat Rev Genet, 22, 203–215.

114. Marguerat, S. and Bähler, J. (2010) RNA-seq: from technology to biology. Cell. Mol. Life Sci., 67, 569–579.

115. Oliveira, D.S., Rosa, M.T., Vieira, C. and Loreto, E.L.S. (2021) Oxidative and radiation stress induces transposable element transcription in *Drosophila melanogaster*. J of Evolutionary Biology, 34, 628–638.

116. Horváth, V., Merenciano, M. and González, J. (2017) Revisiting the Relationship between Transposable Elements and the Eukaryotic Stress Response. Trends in Genetics, 33, 832–841.

117. Nicolau, M., Picault, N. and Moissiard, G. (2021) The Evolutionary Volte-Face of Transposable Elements: From Harmful Jumping Genes to Major Drivers of Genetic Innovation. Cells, 10, 2952.

118. Hug, N., Longman, D. and Cáceres, J.F. (2016) Mechanism and regulation of the nonsense-mediated decay pathway. Nucleic Acids Res, 44, 1483–1495.

119. Harigaya, Y. and Parker, R. (2010) No-go decay: a quality control mechanism for RNA in translation. WIREs RNA, 1, 132–141.

120. Vasudevan, S., Peltz, S.W. and Wilusz, C.J. (2002) Non-stop decay—a new mRNA surveillance pathway. BioEssays, 24, 785–788.

