## Supplementary Figures for "ChimeraTE: A pipeline to detect chimeric transcripts derived from genes and transposable elements"

**Supplementary Figure 1:** ChimeraTE Mode 1 detection of chimeric reads (CR). **a**) TE-exonized embedded transcript: The exon is artificially divided into two portions, based on the TE’s start and end position. Reads with both mates aligned solely into the TE region (gray block) are discarded. Concordant *end-to-end* reads are considered as CRs if one mate is aligned into the TE, and the other is aligned into the exon; while *split* reads are counted when a partial region of one mate is aligned into the TE (or exon) and the other is aligned into the exon (or TE). Such approach avoids counting autonomous TE expression as evidence for chimeric transcripts. **b**) Similarly, TE-exonized overlapped does not consider reads aligned solely to the TE as CR. Instead of it, both concordant and split reads are considered as CRs when both mates have concomitantly aligned into exons and TEs. **c**) CRs supporting TE-exonized intronic are computed when both mates are aligned between the TE and an exon. **d**) CRs for TE-initiated transcripts must have one mate, concordant or split, aligned into the TE located 3kb upstream to the gene, whereas the other is aligned into the exon. **e**) CRs for TE-terminated transcripts have the same rationale as TE-initiated, but with the TE insertion located 3kb downstream to the gene. Any exon can have CRs, even though they are not represented in the figure.

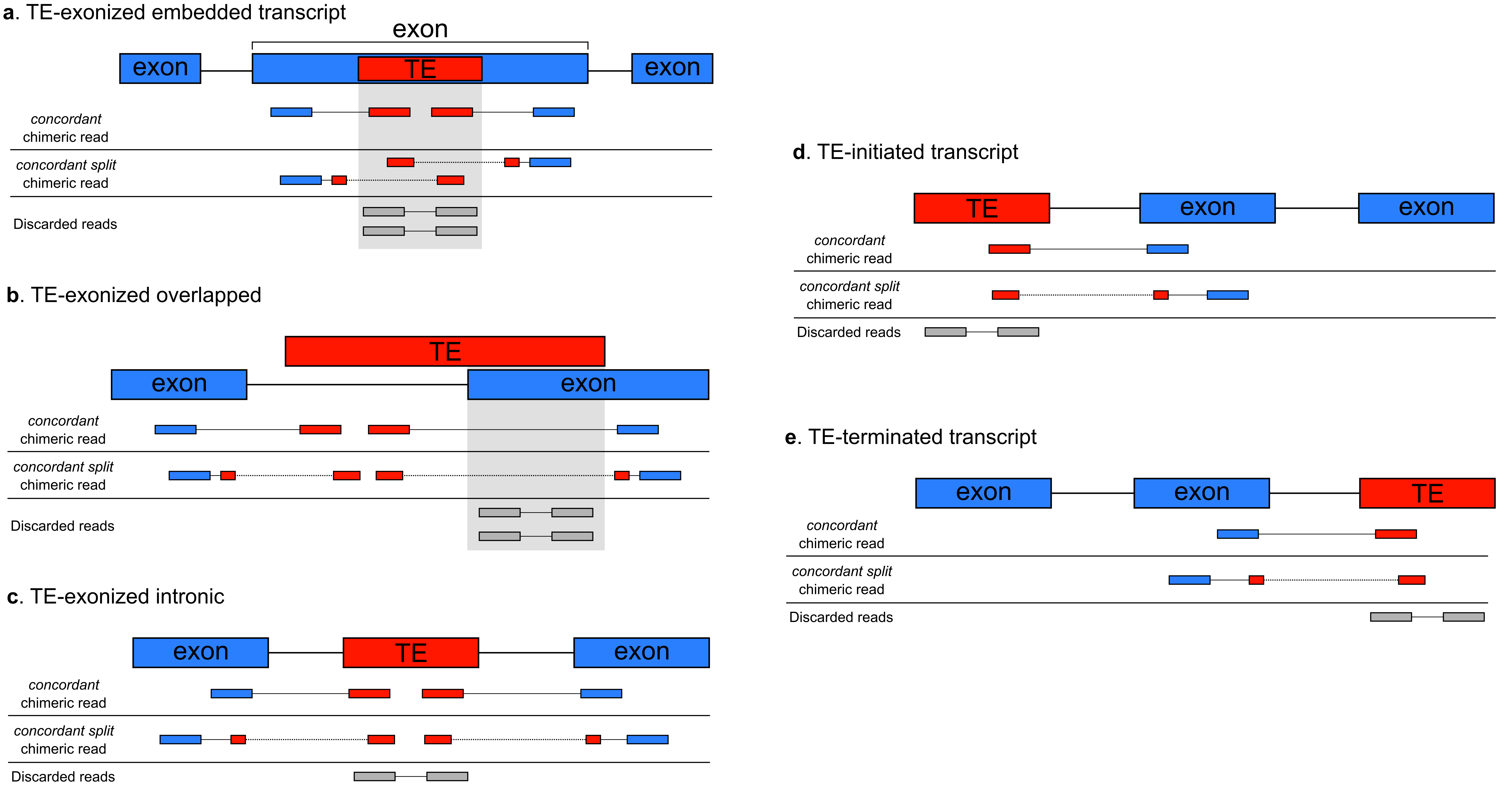

**Supplementary Figure 2**: Frequency of TE copies by TE family that were filtered out after removal of insertions with SSRs covering more than 50% of their length. *Roo* family comprise more than 85% in the four wild-type strains.

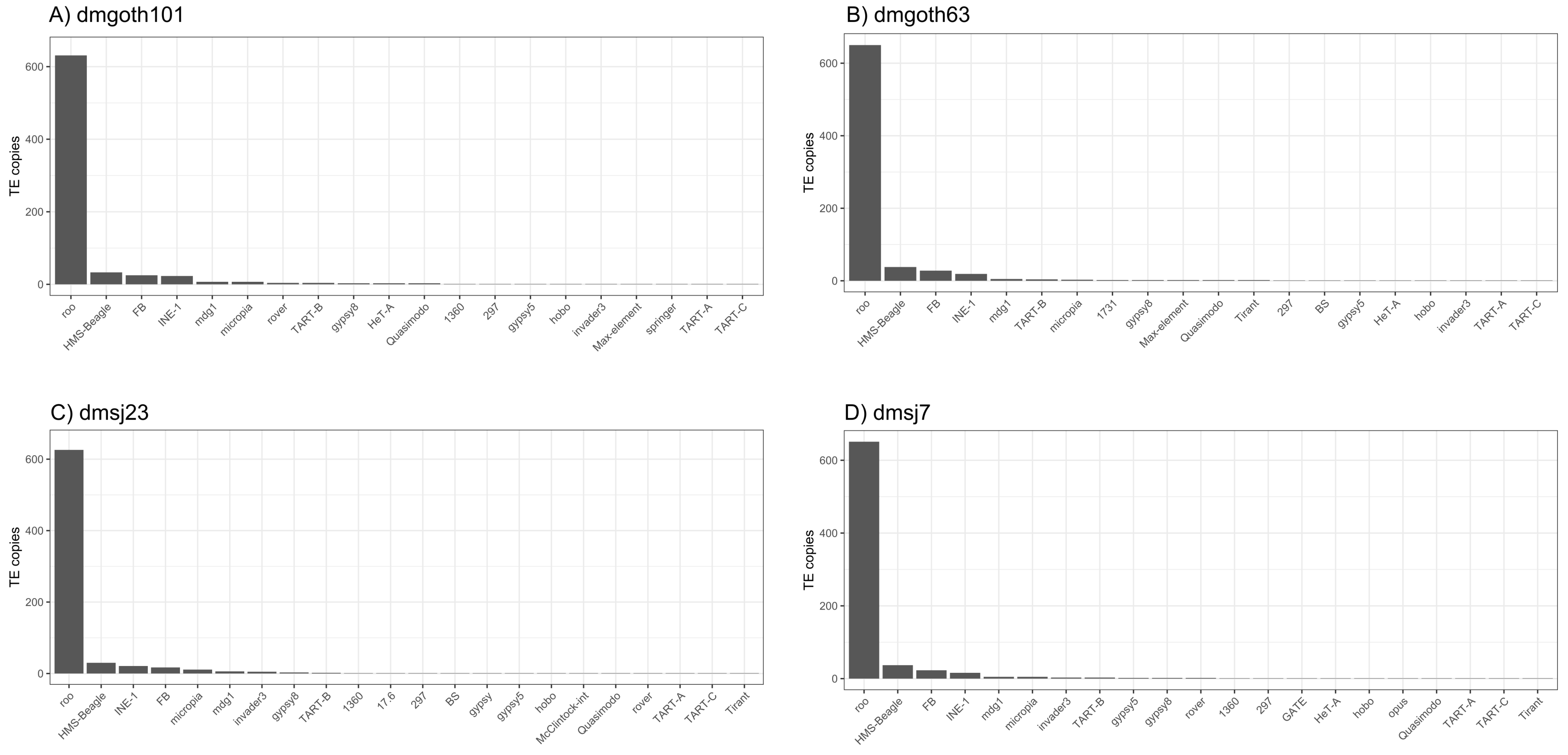

**Supplementary Figure 3**: Average length of TE insertions removed after applying filtering of SSRs > 50% of TE insertion length.

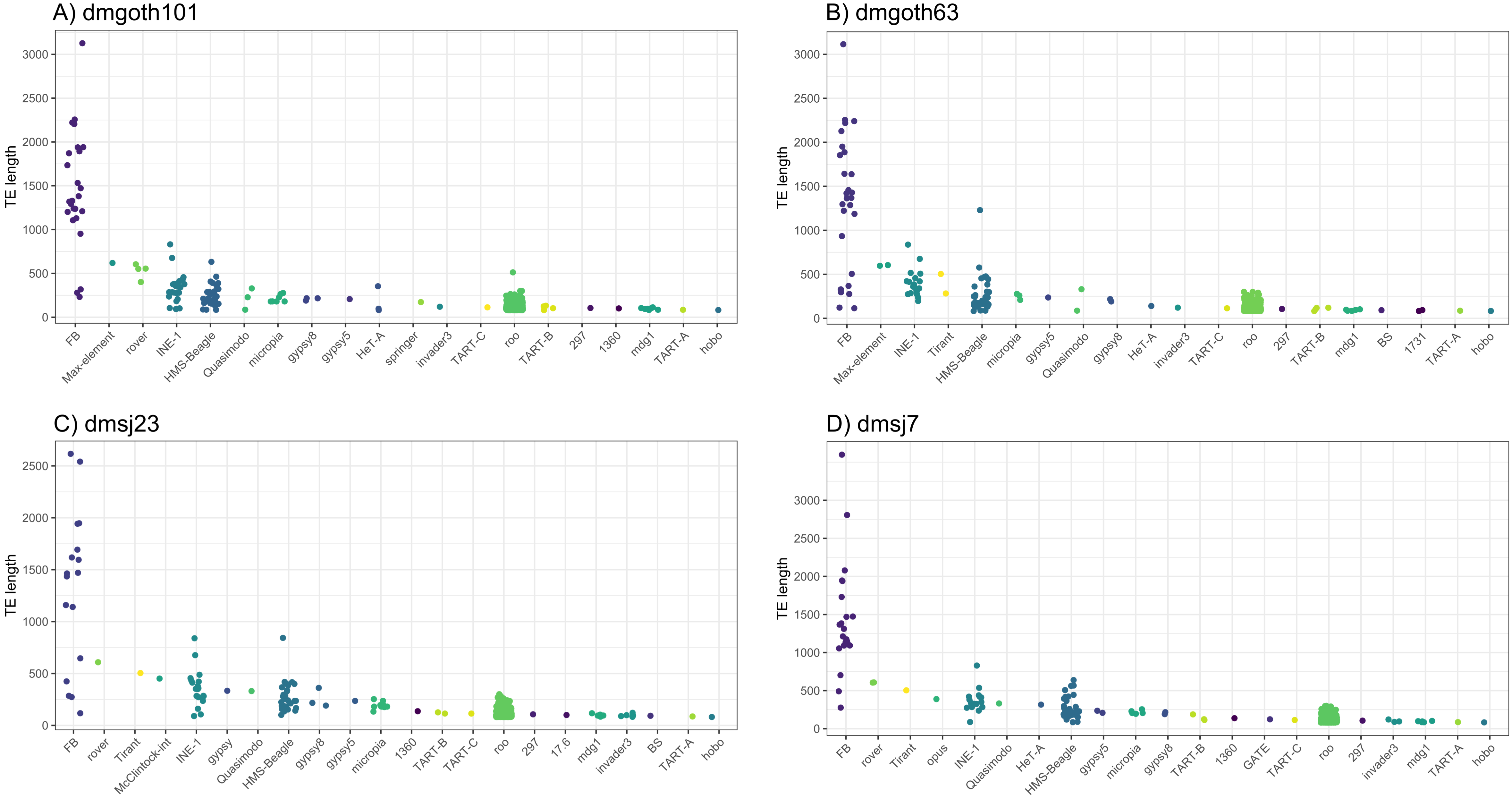

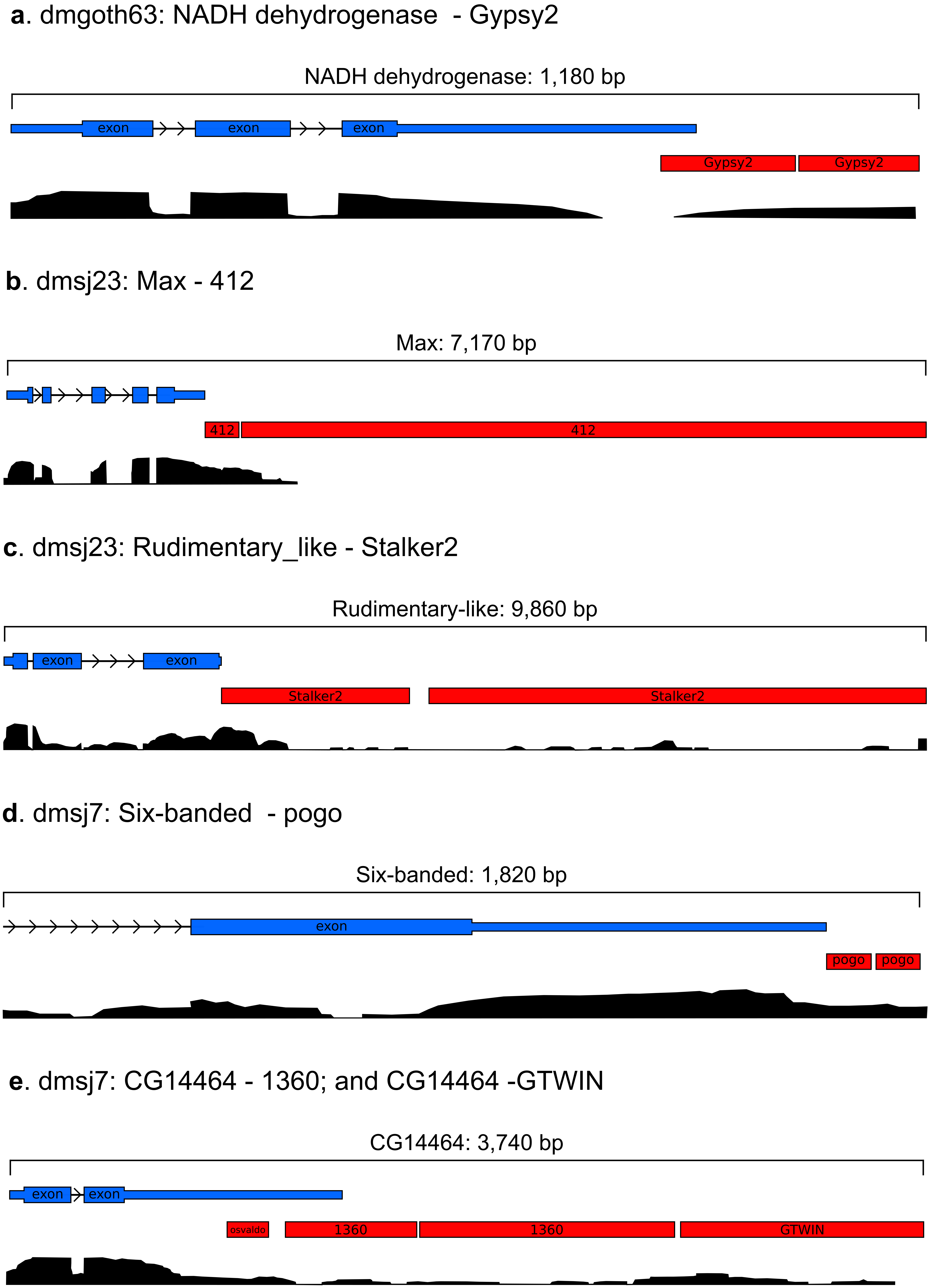
**Supplementary Figure 4**: TE-terminated transcripts with multiple TE insertions.

**Supplementary Figure 5**: Spearman correlation between the number of TEs within genes with FPKM > 1. Y axis represents the number of TE insertions identified as chimeric transcripts, whereas X axis represents the number of TE insertion within gene region.

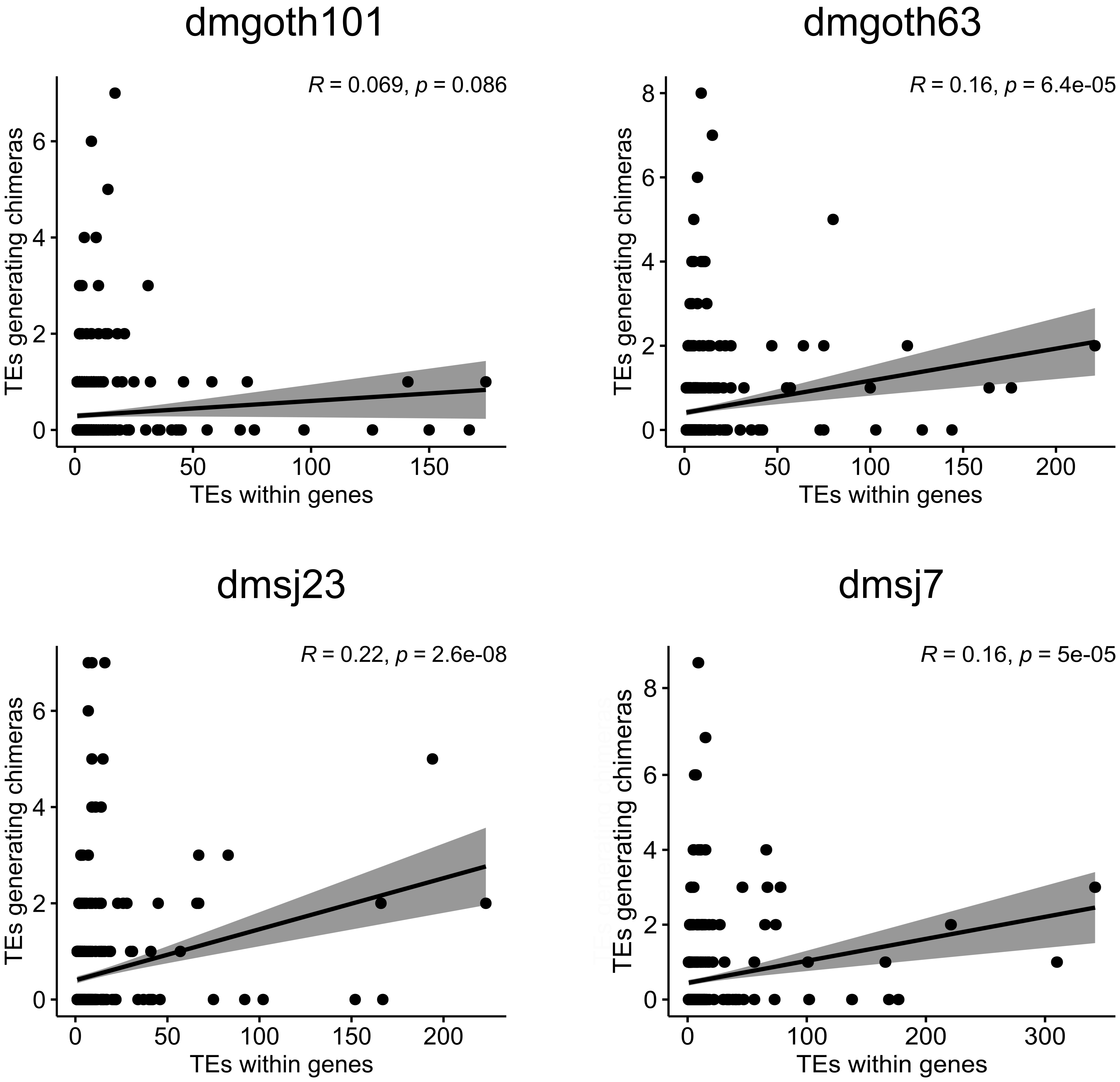

**Supplementary Figure 6**: Spearman correlation between the number of TE insertion inside genes, per TE family, and its respective number of chimeric transcripts.

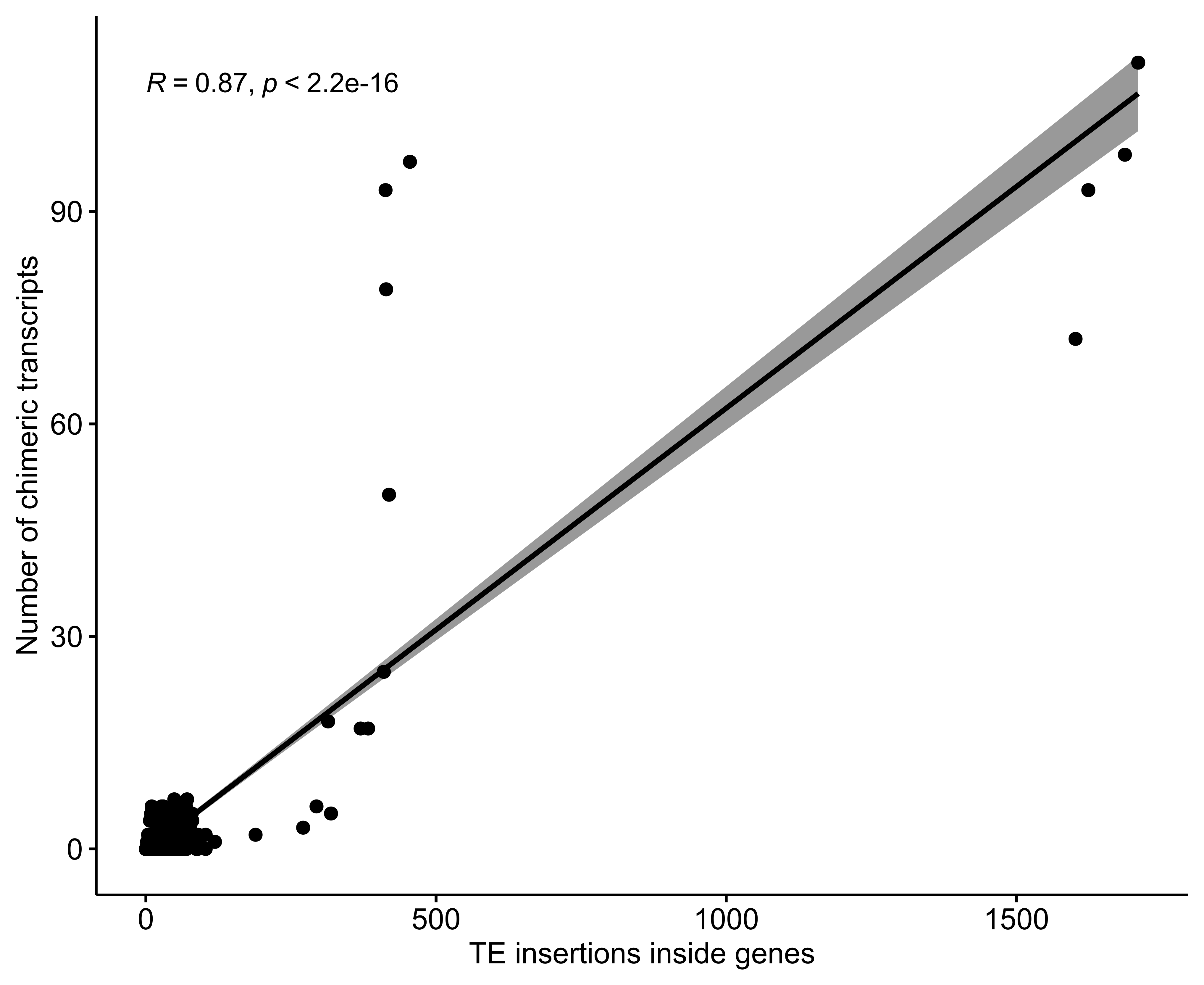

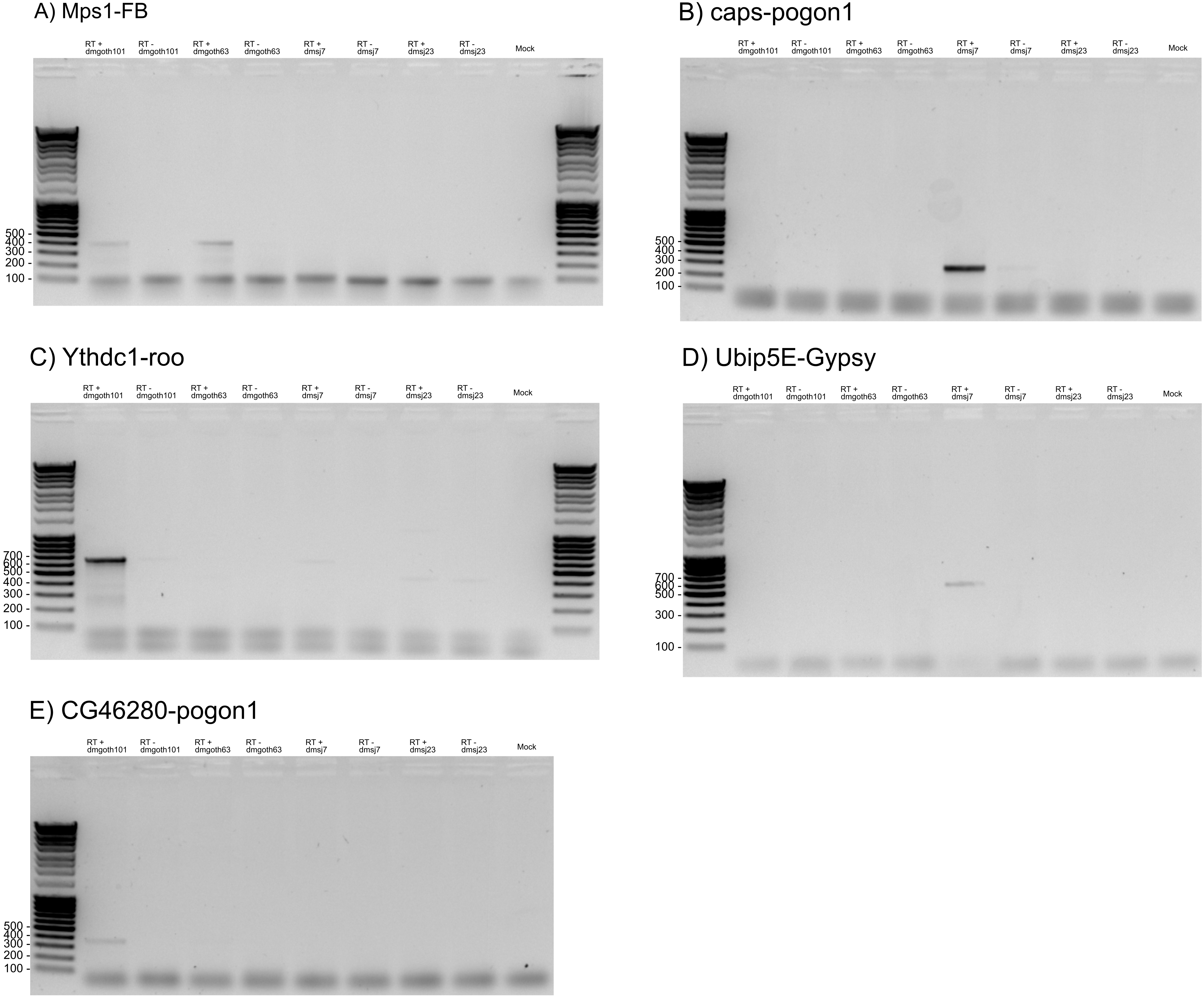
**Supplementary Figure 7**: Five chimeric transcripts found by ChimeraTE Mode 2 in our four wild-type strains derived from TE insertions absent in the reference genome. All RT-PCR amplifications represent the same result obtained by ChimeraTE Mode 2.

**Supplementary figure 8**: **A**) Presence of chimeric transcript derived from *r1* gene and *Stalker2* TE in the four wild-type strains. The RT-PCR reveals amplification in the expected length in both dmsj7 and dmsj23, whereas for dmgoth101 and dmgoth63 there are off target amplification. **B**) PCR result for the Stalker2 insertion, demonstrating the presence of such insertion in both Brazilian strains.

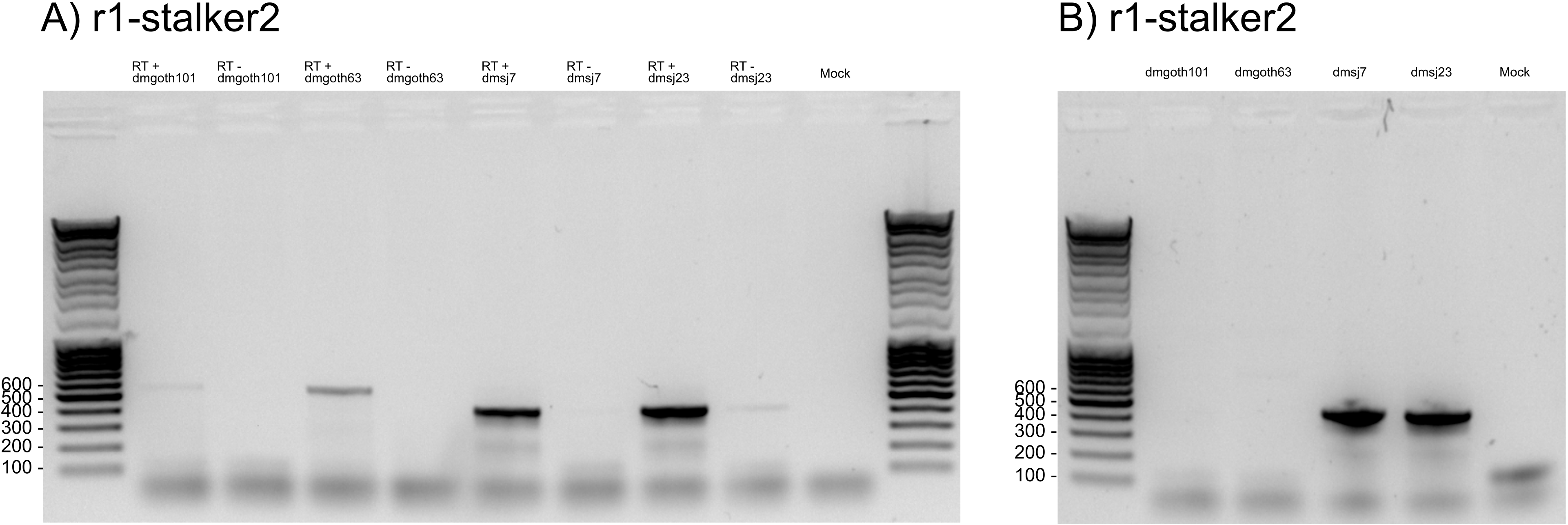

**Supplementary figure 9**: Presence of chimeric transcript derived from *rb* gene and *mdg3* TE in the four wild-type strains.

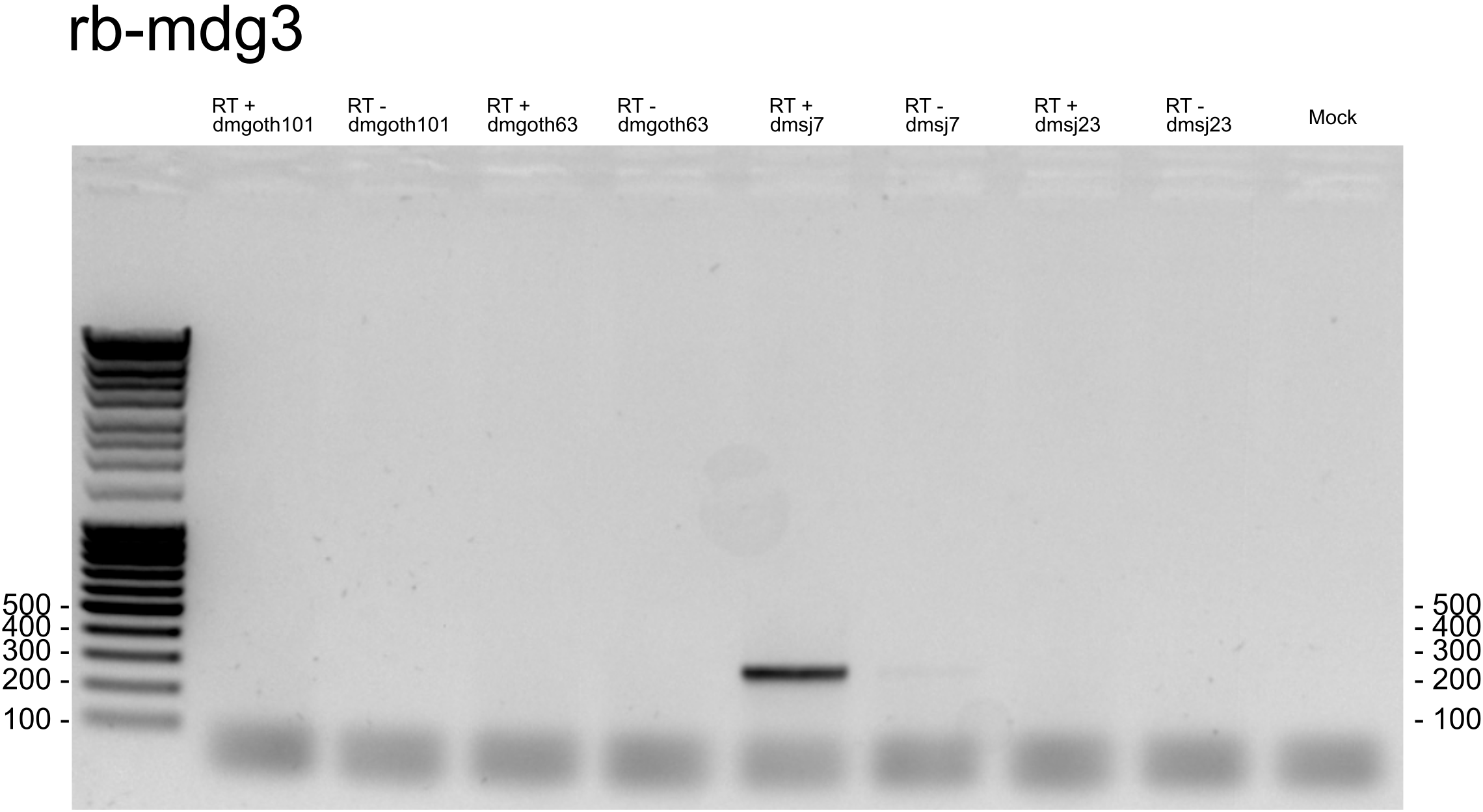

**Supplementary figure 10**: Presence of chimeric transcript derived from *CG1358* gene and *S* TE in the four wild-type strains.

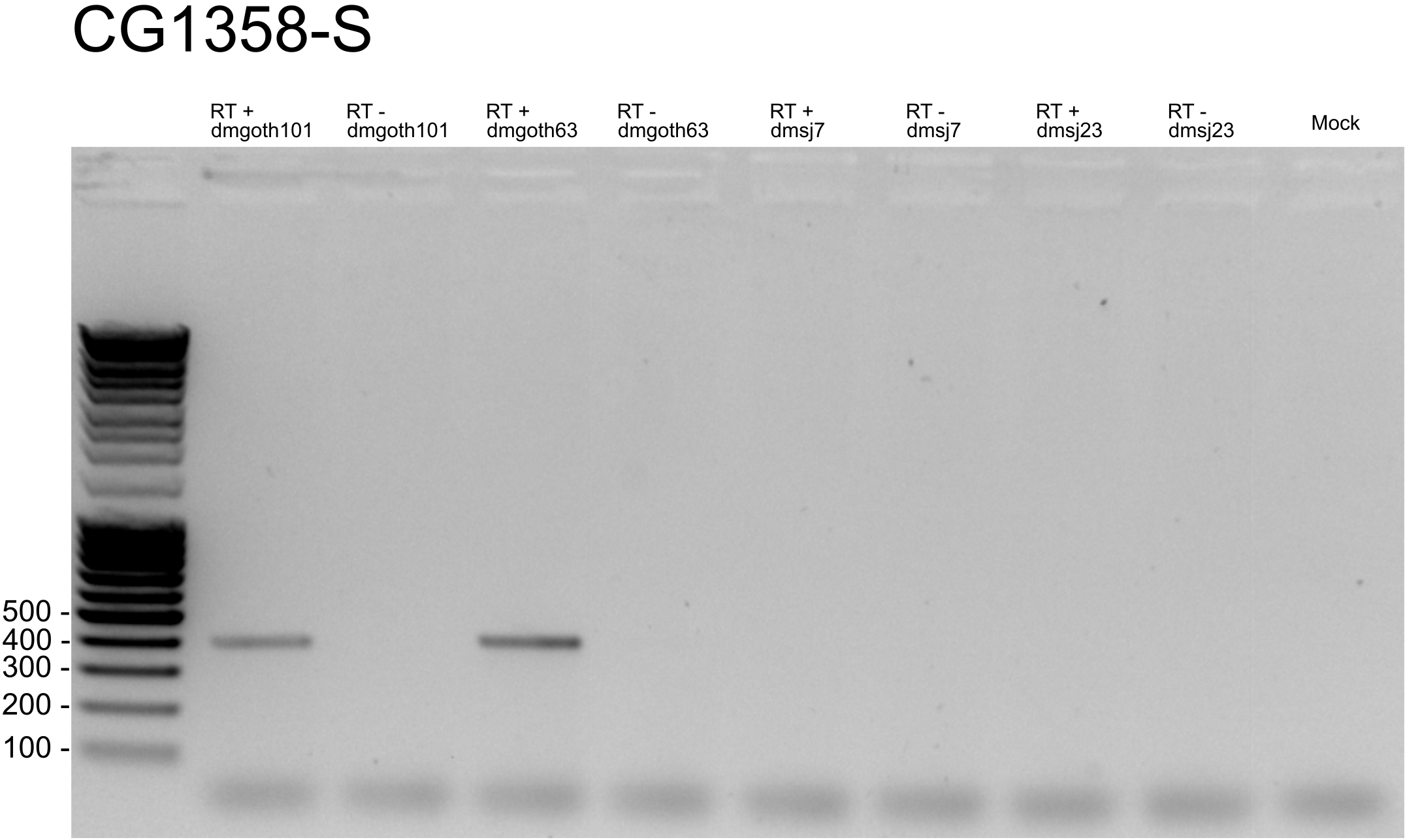

**Supplementary figure 11**: Presence of chimeric transcript derived from *cic* gene and *pogon1* TE in the four wild-type strains.

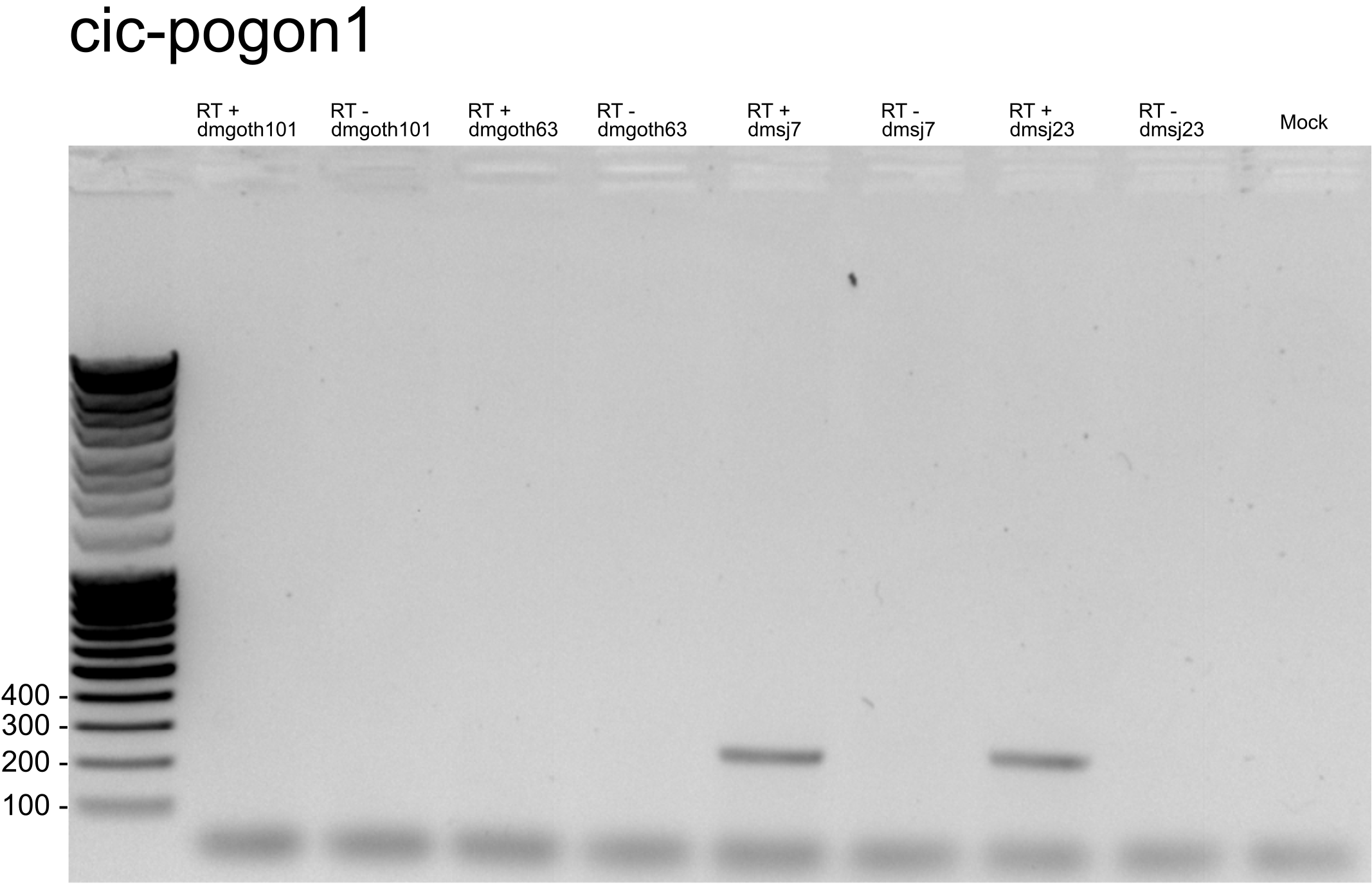

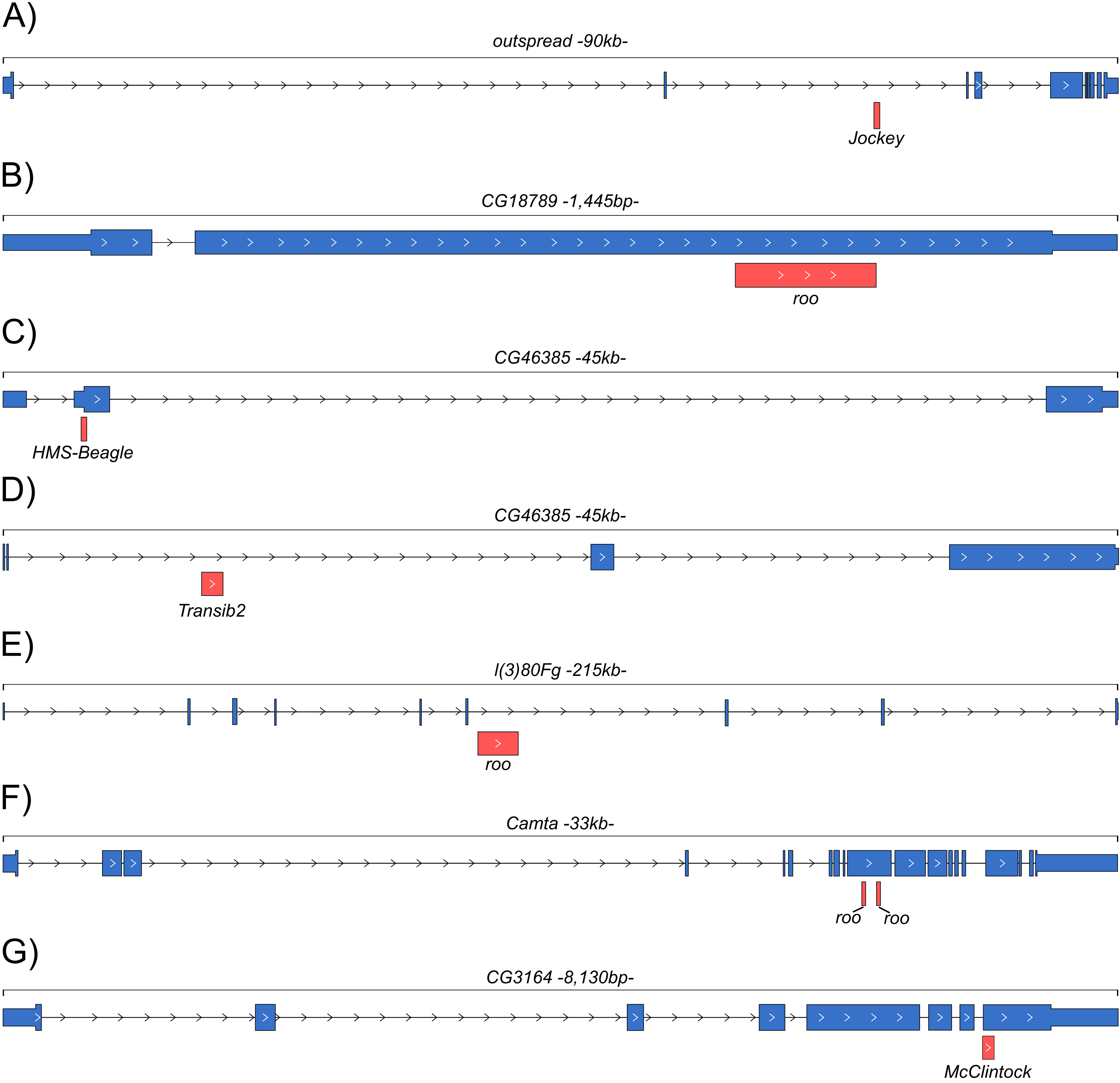
**Supplementary Figure 12**: Gene and TE annotation from *dm6* genome for chimeric transcripts identified by Mode 2 corresponding to genes that were not annotated in the wild-type genome assemblies. Blue boxes: UTRs and CDSs; Red boxes: TE insertion identified by ChimeraTE as the TE family generating the chimeric transcript.

**Supplementary Figure 13**: The total number of TE-aligned reads between both ChimeraTE Modes, in all strains and their respective replicates.

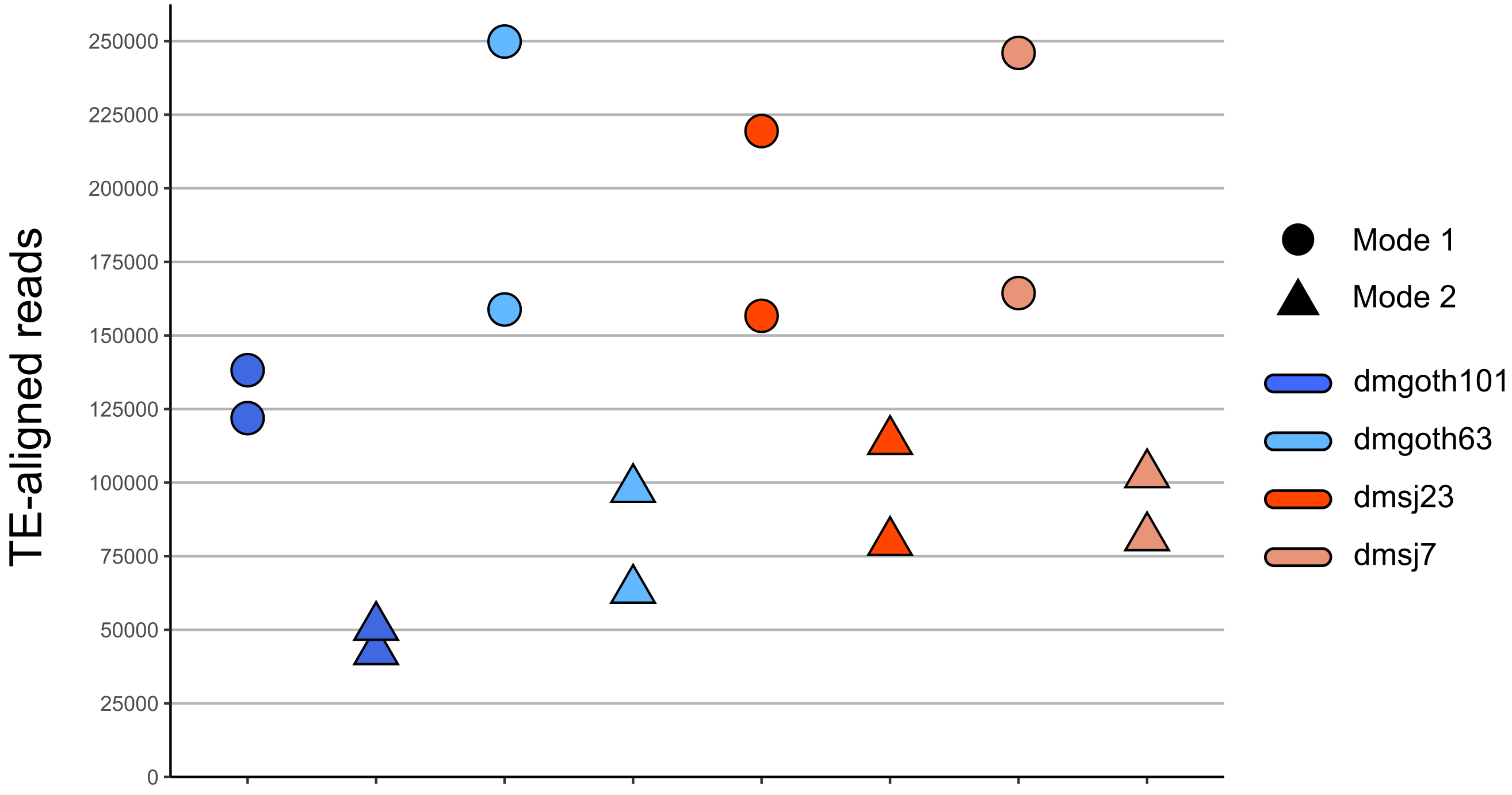

**Supplementary Figure 14**: Positive Person correlations between: A) RNA-seq libraries size and the total of chimeric reads detected in Mode 1 (top) and Mode 2 (bottom); B) TE-aligned reads and the total of chimeric reads detected in in Mode 1 (top) and Mode 2 (bottom).

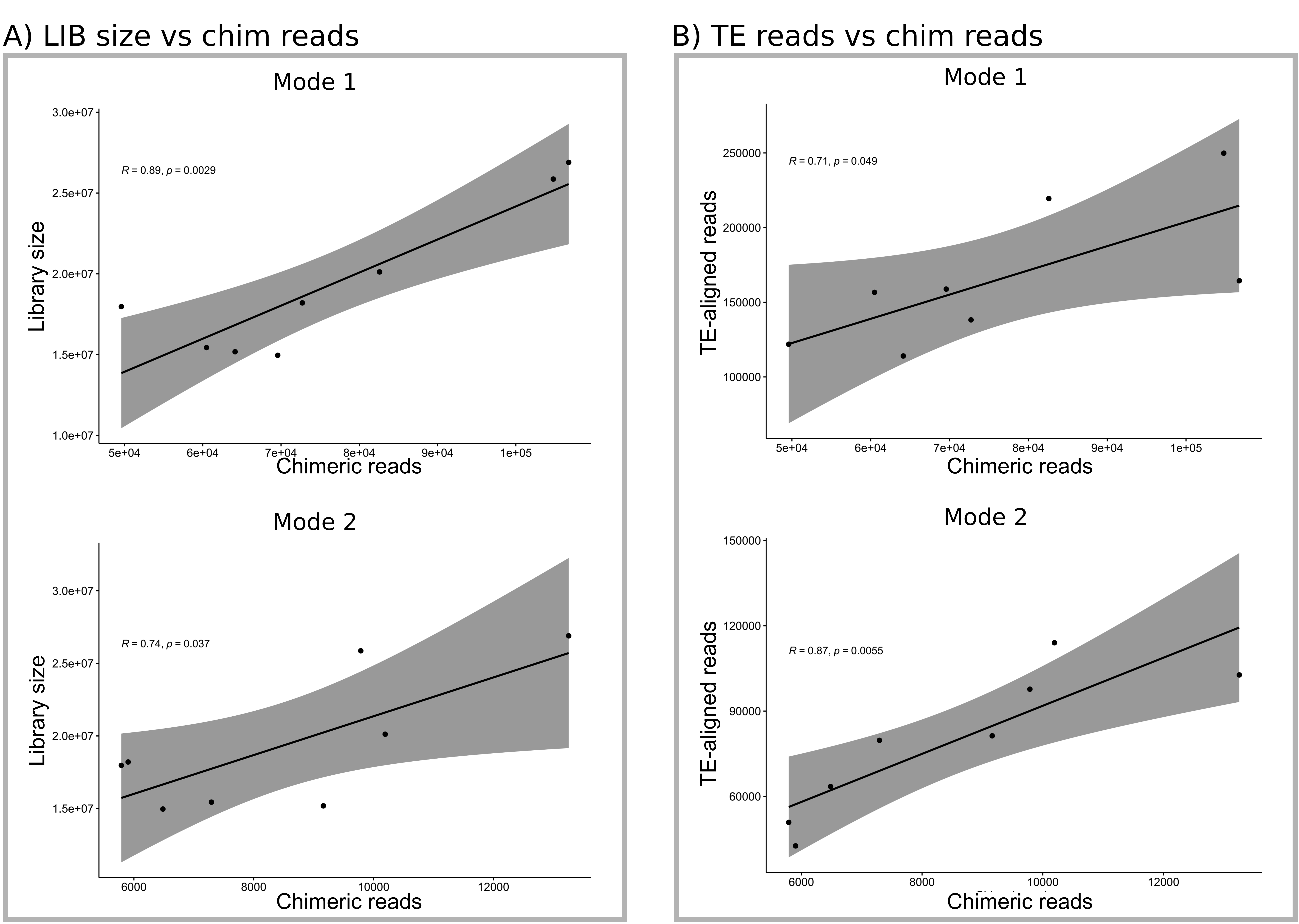

**Supplementary Figure 15**: Chimeric transcripts with TE-derived protein domain found with RNA-seq ONT data.

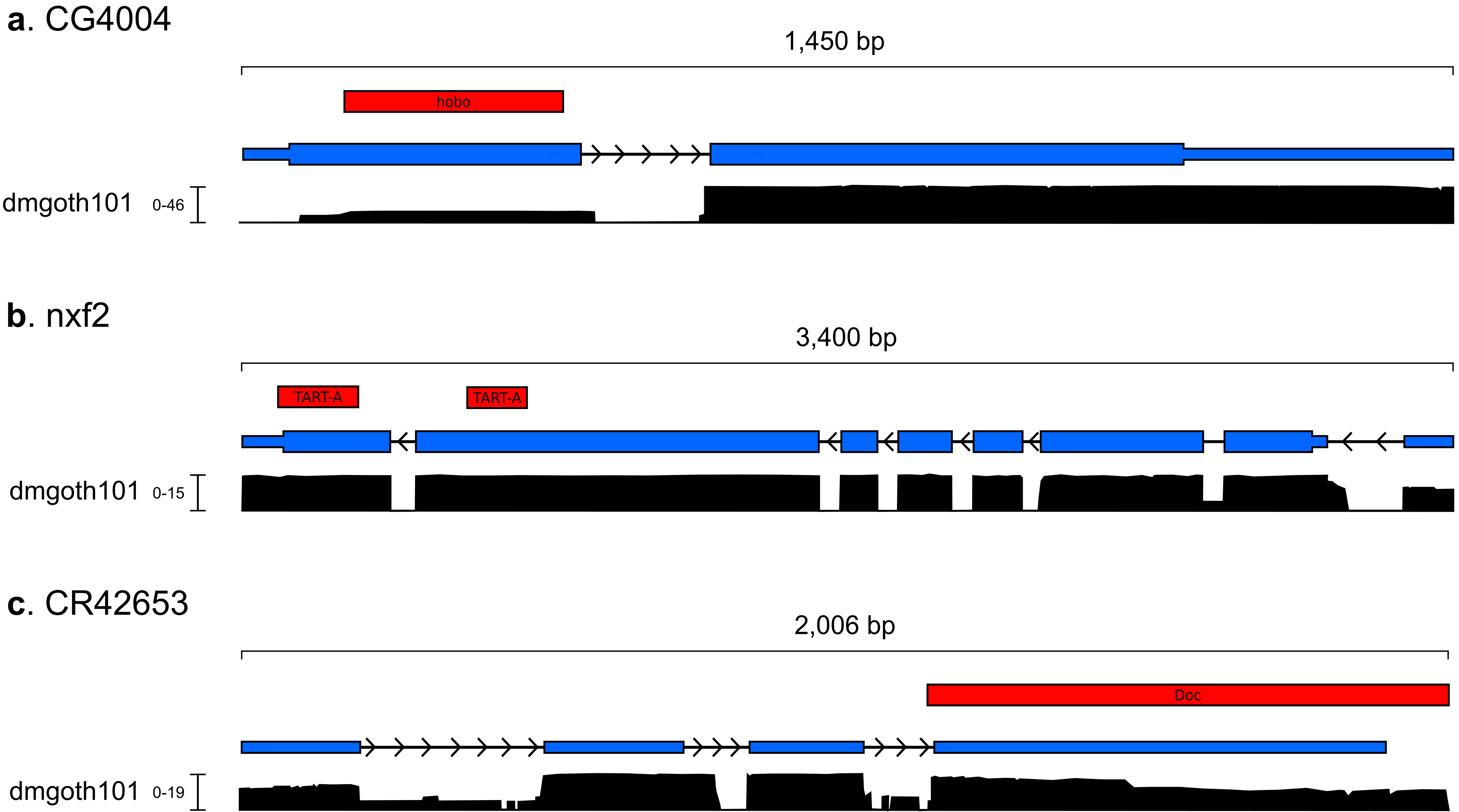

**Supplementary Figure 16**: Relative contribution of TE expression to the gene expression, for genes that are inactive in one strain, but generating chimeric transcript in the other.

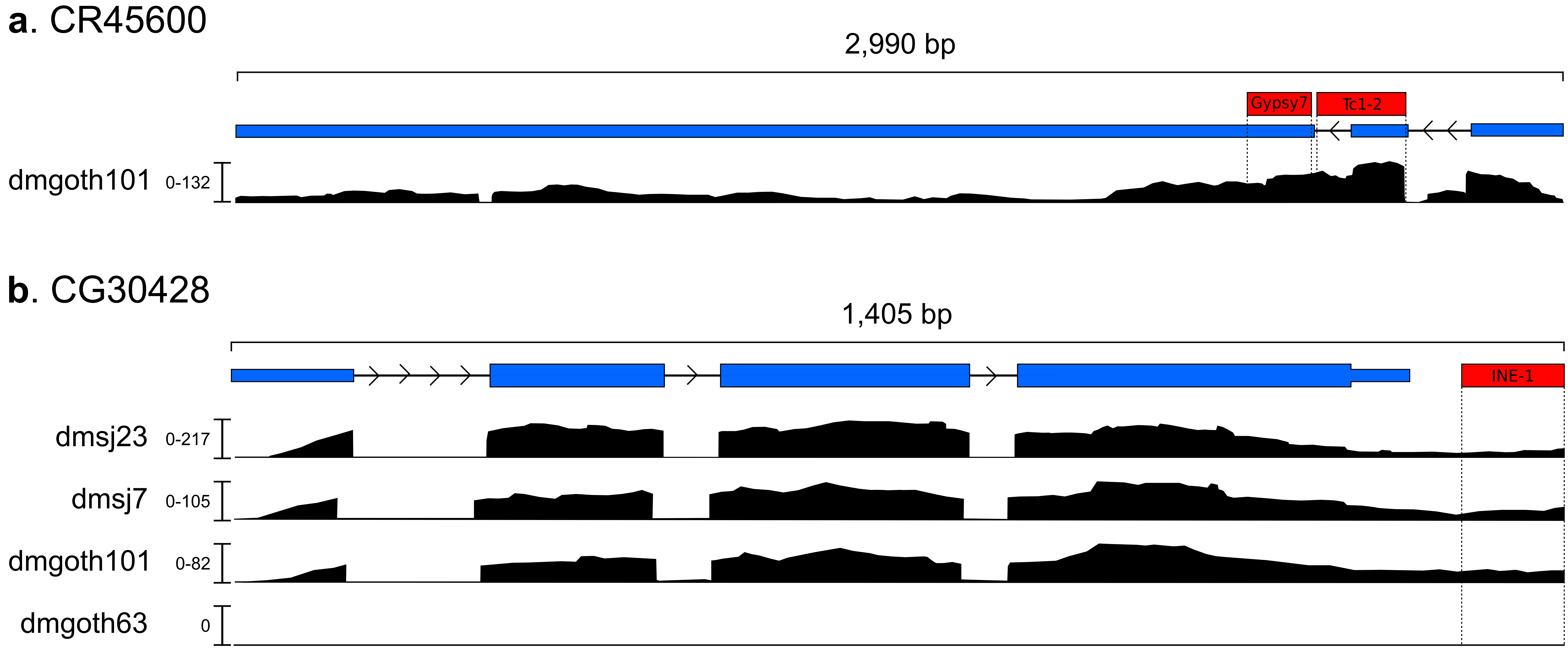

**Supplementary Figure 17**: Correlation between processing-time of ChimeraTE and genome size of species. **a)** Mode 1 has a strong positive correlation with the genome size, since it is the genome-guided Mode, whereas **b)** Mode 2 does not have correlation, most likely due to the de novo transcriptome assembly, which the processing-time is dependent on the library size.

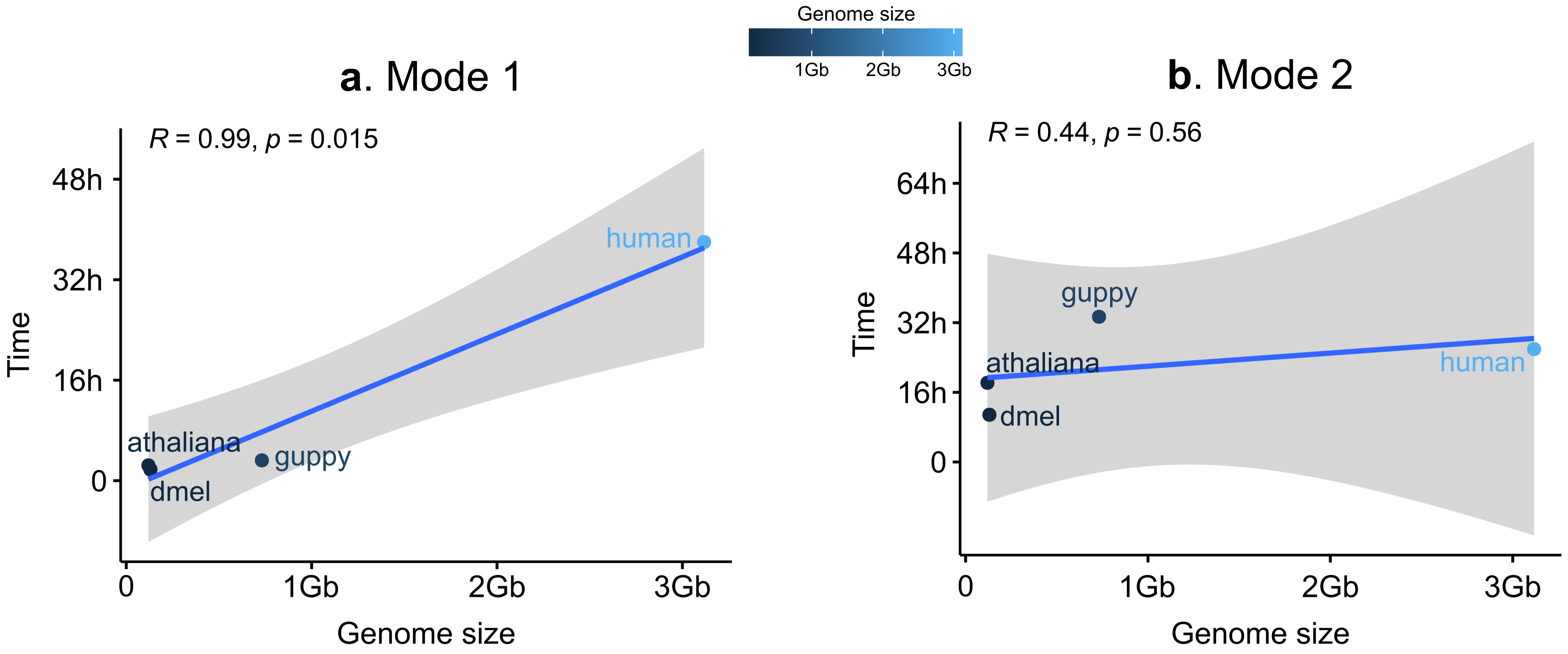
